# Climate and mountains shaped human ancestral genetic lineages

**DOI:** 10.1101/2021.07.13.452067

**Authors:** Pierpaolo Maisano Delser, Mario Krapp, Robert Beyer, Eppie R Jones, Eleanor F Miller, Anahit Hovhannisyan, Michelle Parker, Veronika Siska, Maria Teresa Vizzari, Elizabeth J. Pearmain, Ivan Imaz-Rosshandler, Michela Leonardi, Gian Luigi Somma, Jason Hodgson, Eirlys Tysall, Zhe Xue, Lara Cassidy, Daniel G Bradley, Anders Eriksson, Andrea Manica

## Abstract

Extensive sequencing of modern and ancient human genomes has revealed that contemporary populations can be explained as the result of recent mixing of a few distinct ancestral genetic lineages^1^. But the small number of aDNA samples that predate the Last Glacial Maximum means that the origins of these lineages are not well understood. Here, we circumvent the limited sampling by modelling explicitly the effect of climatic changes and terrain on population demography and migrations through time and space, and show that these factors are sufficient to explain the divergence among ancestral lineages. Our reconstructions show that the sharp separation between African and Eurasian lineages is a consequence of only a few limited periods of connectivity through the arid Arabian peninsula, which acted as the gate out of the Arican continent. The subsequent spread across Eurasia was then mostly shaped by mountain ranges, and to a lesser extent deserts, leading to the split of European and Asians, and the further diversification of these two groups. A high tolerance to cold climates allowed the persistence at high latitudes even during the Last Glacial Maximum, maintaining a pocket in Beringia that led to the later, rapid colonisation of the Americas. The advent of food production was associated with an increase in movement^2^, but mountains and climate have been shown to still play a major role even in this latter period^3,4^, affecting the mixing of the ancestral lineages that we have shown to be shaped by those two factors in the first place.

## Main Text

Recent large-scale analyses of modern and ancient genomes have revealed that most contemporary Out-of-Africa human populations formed during the Holocene as the result of mixing of a limited number of genetically distinct ancestral lineages^1^ (see Fig. 1 for a list of the main lineages). But the origins of those lineages are less clear. The few very early human ancient genomes (e.g. Sunghir^5^, Ust’Ishim^6^) are relatively undifferentiated, and whilst they provide a timing for the split of the Asian and European lineages, they say little about the circumstances that promoted the subsequent separation into the ancestral lineages that contributed to modern populations. The relatively few ancient genomes that predate the Last Glacial Maximum are too sparse to provide a clear picture of these dynamics, and by the time sampling of ancient DNA becomes more extensive (i.e. in the last 10k years), those ancestral ancestral lineages are already well established.

**Fig. 1:**
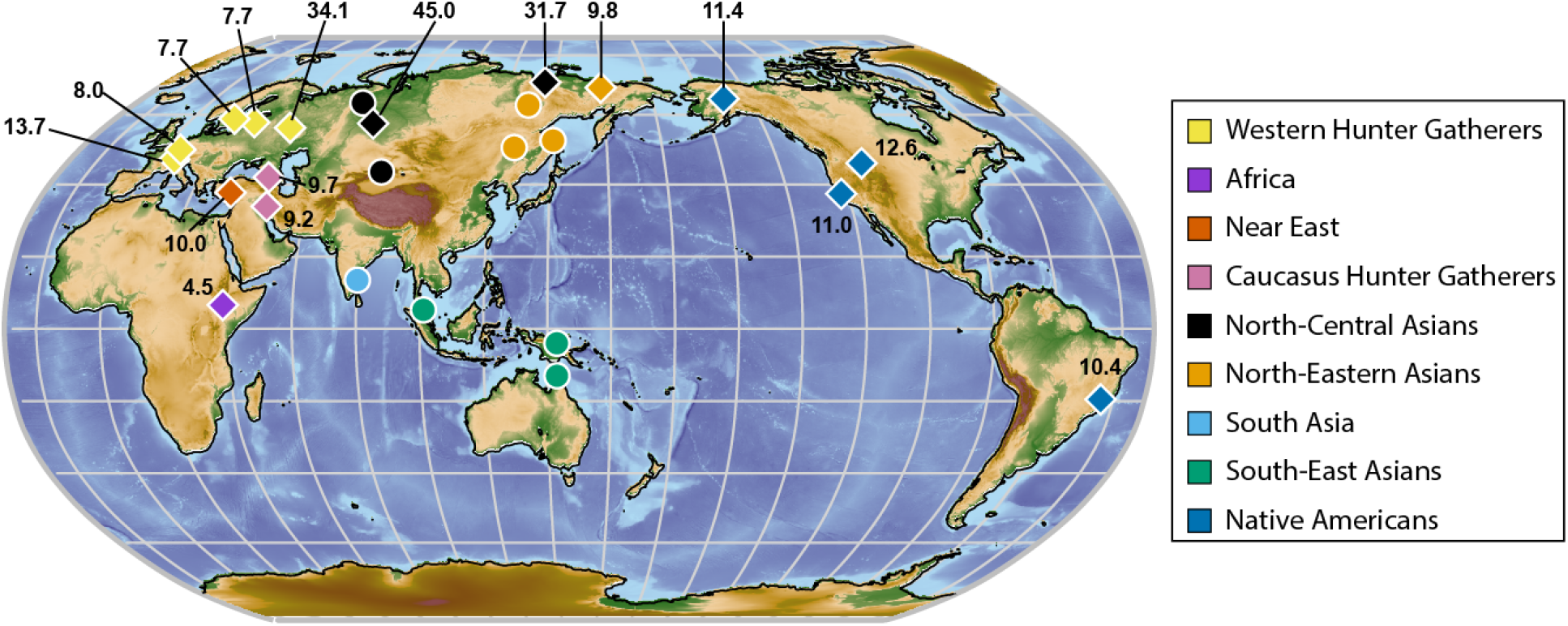
Distribution on ancient (diamonds) and modern (circles) HG genomes. Colours represent the assignment to ancestral lineages. Individual Yana1 (dated at 31.7 kya) showed a greater genetic affinity with samples from North-Central Asia than North-Eastern Asia; therefore it has been included in the former ancestral lineage group (**Extended Data Fig. 1**). Numbers represent the mean date in thousands years BP for ancient samples.

So, what processes might have promoted the divergence of the ancestral human lineages? Climate and terrain are often invoked as major determinants of the degree of movement over the landscape, and indeed their signature can still be found in the level of genetic differentiation among contemporary populations despite high movement rates in historical times^3^. It is thus plausible that climate and terrain played a role in the emergence of the ancestral human lineages that have been identified out of Africa (poor ancient DNA preservation means that we only have a limited understanding of African ancestral lineages, so we will not investigate them in this paper). Whilst the role of climate in determining the timing of the Out-of-Africa has received a great deal of attention^7–9^, quantifying its role in the routes and dynamics of the spread has been much more challenging due to the limited archaeological record from the early part of the expansion.

### Climate and mountains are sufficient to explain divergences among lineages

In this paper, we formally test the role of climate and terrain in shaping the genetic makeup of humans out of Africa by modelling the genetic divergence of a panel of high coverage modern and ancient hunter-gatherer (HGs) genomes representing the main ancestral lineages (**Fig. 1, Extended Data Table 1 and 2**). We focus on HGs as all ancestral lineages arose before the advent of food production, which on the other hand is associated with large scale movements that led to their mixing. Specifically, we used climate informed spatial genetic models (a further development of ^9^) that reproduce the world as a lattice of hexagonal cells, with coastlines changing through time according to sea level changes. The demography within each cell depends on reconstructions of local, time-varying climatic conditions of the past (see Methods). We explored a large number of values for the parameters that link climatic variables and mountains to local effective population sizes and migration rates (see **Extended Data Table 3 and 4**), and selected the values that give the best match to the pattern of genetic pairwise differentiation (π_between_) between ancestral lineages using an Approximate Bayesian Computation framework.

The effects of climate and mountains were sufficient to explain the divergence among ancestral human lineages out of Africa. The model was able to reconstruct all pairwise divergences simultaneously, as seen by inspecting pairwise plots of divergences among groups (**Extended Data Fig. 2**). Furthermore, we tested the goodness of fit by comparing the median of Euclidean distances between observed π_between_ and the estimates for the best set of 2% (n=5058) simulations, to the distribution of Euclidean distances between randomly sampled simulations (1000 replicates) and their respective best sets. For a model that can replicate the observed statistics, the median of Euclidean distances of the observed data should not differ significantly from the distances of random simulations, and this is indeed the case for our model, which had a p-value of 0.465 (**Extended Data Fig. 3**). A separate line of validation of the realism of our model comes from comparing the distribution of our estimated effective population sizes based on genetics (N_e_) to that of census sizes from ethnographic surveys (N_census_). N_e_ represents the idealised size of a randomly mating population, so we do not expect a 1:1 relationship between these two quantities; however, we would expect a good match in their spatial distribution, and indeed we find a high and significant spatial congruence (Pearson’s r corrected for spatial autocorrelation: 0.61; p<0.001), with similar regions of high and low densities (**Extended Data Fig. 4**).

### Aridity explains the split between African and Eurasian lineages

The timing of the Out-of-Africa migration was linked to a suitably wet period to allow the exit through the nowadays mainly arid Arabian Peninsula. In our simulations, we constrained the model to prevent an exit out of Africa before 70k years ago. Whilst archaeological evidence has demonstrated human expansions prior to this point^10^, made possible by climatically favourable windows, early colonists failed to permanently settle in Eurasia in significant numbers (possible due to competition with other hominins^7,8^ which our model does not account for). Indeed, all Out-of-Africa populations have a divergence time that imply a more recent exit^3^ (with the exception of a possible very small contribution of an early wave into Papuans, but see^11^), and we therefore did not explore the scenario of an earlier exit. Given this constraint, the precipitation threshold needed by humans to persist and expand was estimated between ∼102 and 115 mm/yr (95% credible interval)(**Fig. 2a**) to obtain good fits to the genetic patterns. This threshold makes ecological sense, as it is approximately the required amount of annual rainfall before a desert turns into a xeric shrubland^12^. Indeed, a similar threshold is found when looking at where contemporary HGs live, as well predicting the presence of grazers in the animal community^8^. A threshold above 120mm/yr of rainfall would have prevented any exit until the wet Holocene (**Fig. 2b**). The lower threshold values selected by the model (102-115mm/yr) allowed an exit ∼60kya, with some intermittent connectivity for the following 30 kyrs (**Fig. 2b**). However, the model strongly rejected thresholds lower than 100mm/yr, which would have allowed, over the period from 60 to 30 kya, for extensive connectivity between Africa and populations close to the exit point, such as those in Anatolia and the Zagros Mountains (the Near East lineage and the southern range of the Caucasus HGs). This high input by African lineages into these populations, in turn, would have resulted in an excessive increase in their divergence from other Out-of-Africa populations, as clearly shown when focussing on Western HGs (**Fig. 2c**): only intermediate values of the precipitation threshold lead to the correct amount of divergence between populations in Anatolia and the Zagros Mountains on the one hand and the Western HGs on the other.

**Fig. 2:**
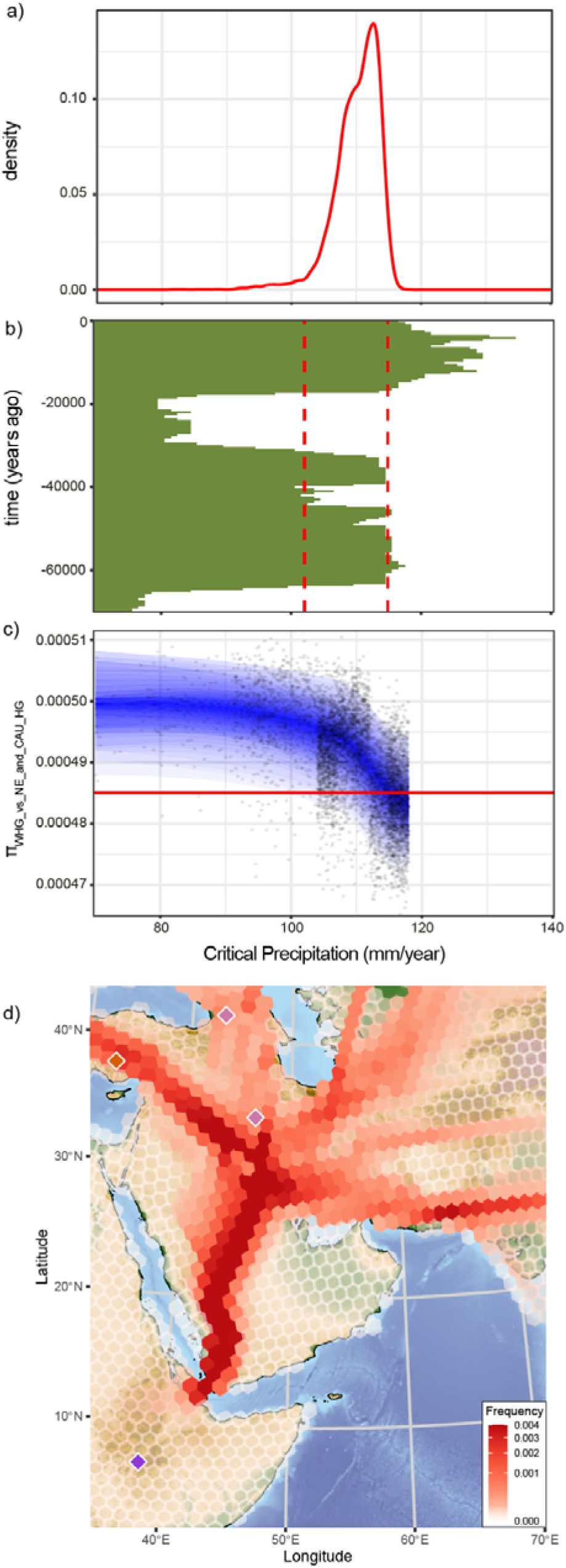
a) The posterior probability for critical precipitation (mm/yr) defining the minimum amount of mean annual precipitation required for population persistence. b) Periods of connectivity between Africa and Eurasia depending on critical precipitation. Red dashed lines represent the 95% credible interval of the posterior distribution for critical precipitation. c) The effect of critical precipitation and the divergence between Western HGs and populations in Anatolia and the Zagros mountains for the 2% of best fitting models used in the ABC. Blue shading shows the 95% interval, with darker shading showing progressively tighter quantiles; the observed divergence is shown by the horizontal red line. d) Distribution of common ancestor events for the last 70 kyrs. Sampled populations are shown with the same colours used in Fig. 1.

Our model tracks the movement of individuals through time and space. Geneflow, the movement that mattered in shaping the genetic makeup of populations, can then be visualised by following the ancestors of different individuals through time, and plotting the location where common ancestors lived. These maps of common ancestors show the key colonisation routes as well as the migratory links that impacted the genetics of populations. When focussing on the distribution of common ancestor events for the last 70k years for Out-of-Africa populations, we found that, in our model, the southern route (through the Bab-el-Mandeb strait) was the main contributor to the expansion, whilst the northern (through the Isthmus of Suez) played little or no role (**Fig. 2d**). In our simulations, the southern route was set as possible when the sea level was at its minimum, and thus the strait could have been mostly easily crossed^13^. A more thorough discussion of the relative crossability of these two routes is provided by^8^; different assumptions could lead to different relative contributions, and the lack of ancient genomes from the appropriate regions prevents us from testing alternative scenarios. Thus, we caution against using our results as evidence for the southern route; what the results indicate clearly is that that connectivity had to be limited and intermittent to generate the appropriate Out-of-Africa bottleneck and the divergence among populations in Eurasia.

### Mountains and deserts shaped divergences within Eurasia

Mountain ranges were major determinants of the colonisation routes taken once out of Africa, and played a key role in promoting the separation among several ancestral lineages. The model favoured intermediate costs of crossing mountains (*α*_*mountain*_) (**Fig. 3a**). The effect of mountains on modulating migration is best seen when considering the divergences of Western HGs and South Asian lineages from Africa (the route between them crosses a number of mountain ranges, thus providing a strong cumulative effect of altitude): realistic levels of genetic divergence could only be obtained with intermediate costs (**Fig. 3b**). The effect of individual mountain ranges is best visualised by considering the location of common ancestor events, thus capturing gene flow through time. The first split was between Asians, which turned East of the Zagros mountains that they encountered as they moved out of Africa, and Europeans, who turned left and partially crossed them (**Fig. 3c**). The Caucasus acted as an important barrier, as it can be seen from the reduced gene flow through this route; indeed, Caucasus HGs, who reside south of the Caucasus, are genetically distinct from the Western HGs found north of this mountain range^14^. Reduced migration over mountain ranges also played a role in generating the divergence between Southern and Northern Asians, with two streams of gene flow that moved north and south of the Himalayas (and the arid Gobi desert); these two streams met in East Asia (**Fig. 3d**). Because of the lack of barriers at higher latitudes, there was contact between the Eastern range of the Western Eurasian HGs and the Northern Asians (**Fig. 3e**, the Urals are not extensive enough to prevent this contact). This matches the mixed ancestry signals found in the Malt’a genome (24,000-year-old individual from south-central Siberia)^15^, which was not included in our analysis due to its low coverage.

**Fig. 3:**
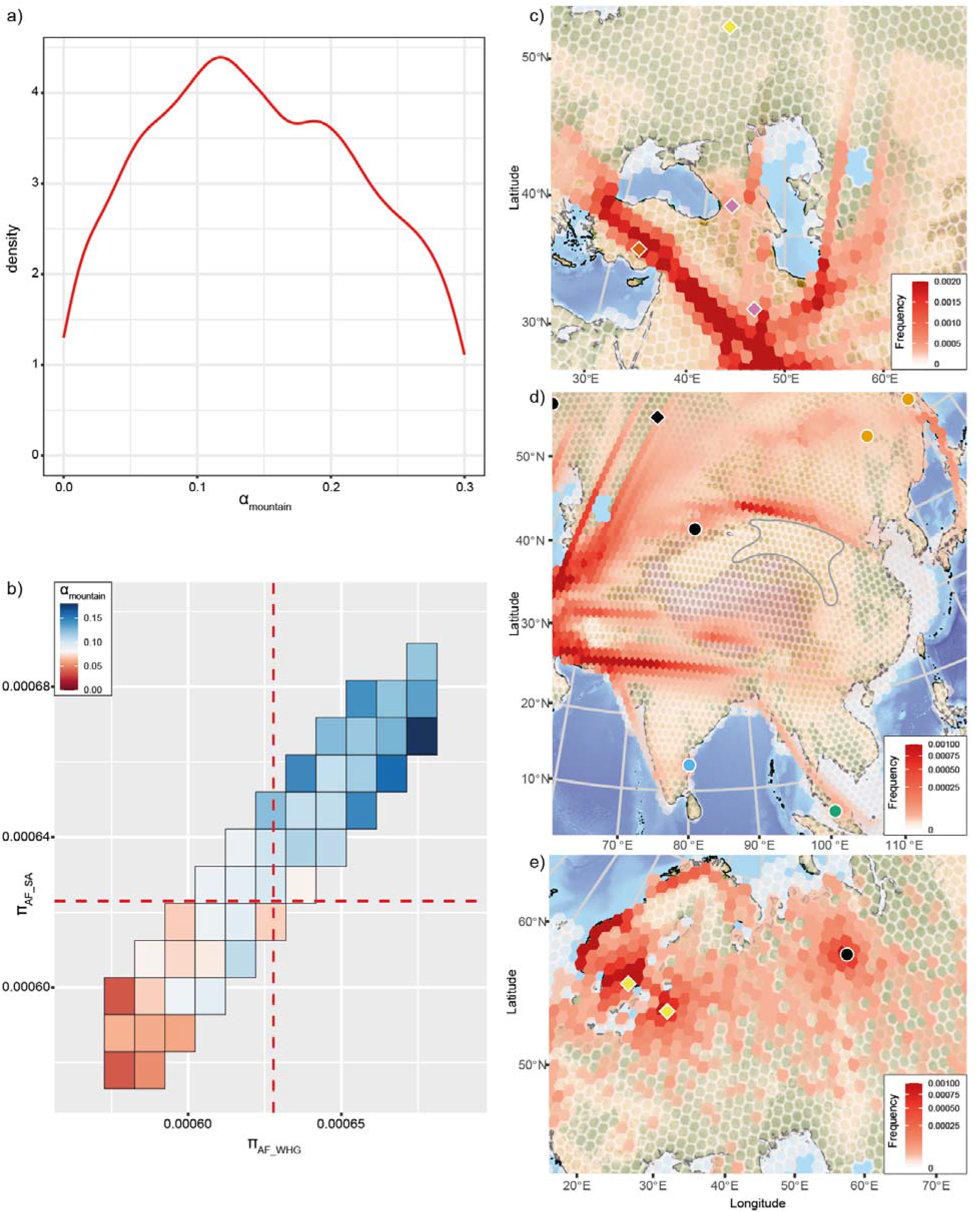
a) Posterior probability for the *α*_*mountain*_ affecting migration between cells. b) The effect of mountain ranges on the divergence between the 4.5kya African sample Mota and Western Hunter Gatherers and South Asian hunter gatherers. African divergence with Western and South Asian hunter gatherers require intermediate values of *α*_*mountain*_ corresponding to the peak of the posterior distribution in panel a). Dashed red lines represent the observed genetic divergence. Distribution of common ancestor events for the c) Caucasus (last 70kyrs), d) Himalayas (last 70kyrs, grey contour represents the Gobi desert) and e) Urals (cropped to the last 12.5 kyrs to emphasise migrations). Sampled populations are shown with the same colours used in Fig. 1.

The southern Asian stream continued into South East Asia, reaching Papua and crossing to Australia (Fig. 3c). The timing of colonisation of Australia is highly dependent on when the Wallace Line can be crossed in the model. As for the Bab-el-Mandeb strait, we took the simple assumption to allow crossings at the lowest sea levels (between 65 and 17 kya); this leads to a somewhat late crossing into Australia at ∼60 kya (**Extended Data Fig. 5**), but we caution that alternative scenarios of an entry several thousands of years earlier cannot be tested due to the lack of ancient genomes from that region.

### Cold temperatures and the colonisation of the Americas

The archeological record suggests human presence at high latitude in very cold environments, as demonstrated by the Yana individual found in Siberia 31 kya (**Fig 1 and Fig. 4a**). In our model, the ability to persist at high latitudes is based on two variables: the optimal temperature (at which humans reach maximum population size) and a temperature tolerance that mediates the cost of living away from that optimum (see Methods). We combined the effect of these two variables for the best fitting scenarios to assess the effect of temperature on the relative population size (i.e. isolating the effect of temperature from the other variables that determine N_e_). The best fitting scenarios all implied an ability to persist in cold environments, such as the area inhabited by the Yana individual; however, population sizes in these regions were very low (**Fig. 4b**). Our model predicted persistence in Beringia up until the end of the Last Glacial Maximum, about 20 kya (**Fig. 4c and d**), thus supporting a Beringian standstill scenario, also supported by other genetic analysis and archaeological evidence^16,17^. In this scenario, the earliest colonists of the Americas originated from this stable pocket close to extensive North American Ice Sheets (**Fig. 4e**). The colonisation started relatively early, between 17 and 16 kya, with the opening of the coastal corridor along the Eastern Pacific coast^18^ (**Fig. 4f**). Recent archaeological work has suggested the possibility of an even earlier arrival in the Americas^19^; our model does not support that, but it should be noted that the population dynamics that we reconstruct are driven by the available genomes. Thus, if there was an earlier colonisation that left little genetic signal in the currently sequenced ancient genomes, we would not select a demography compatible with it. An early colonisation raises the issue of what happened to those early colonists, given that there is no genetic signal of their existence in modern or ancient Native American genomes.

**Fig. 4:**
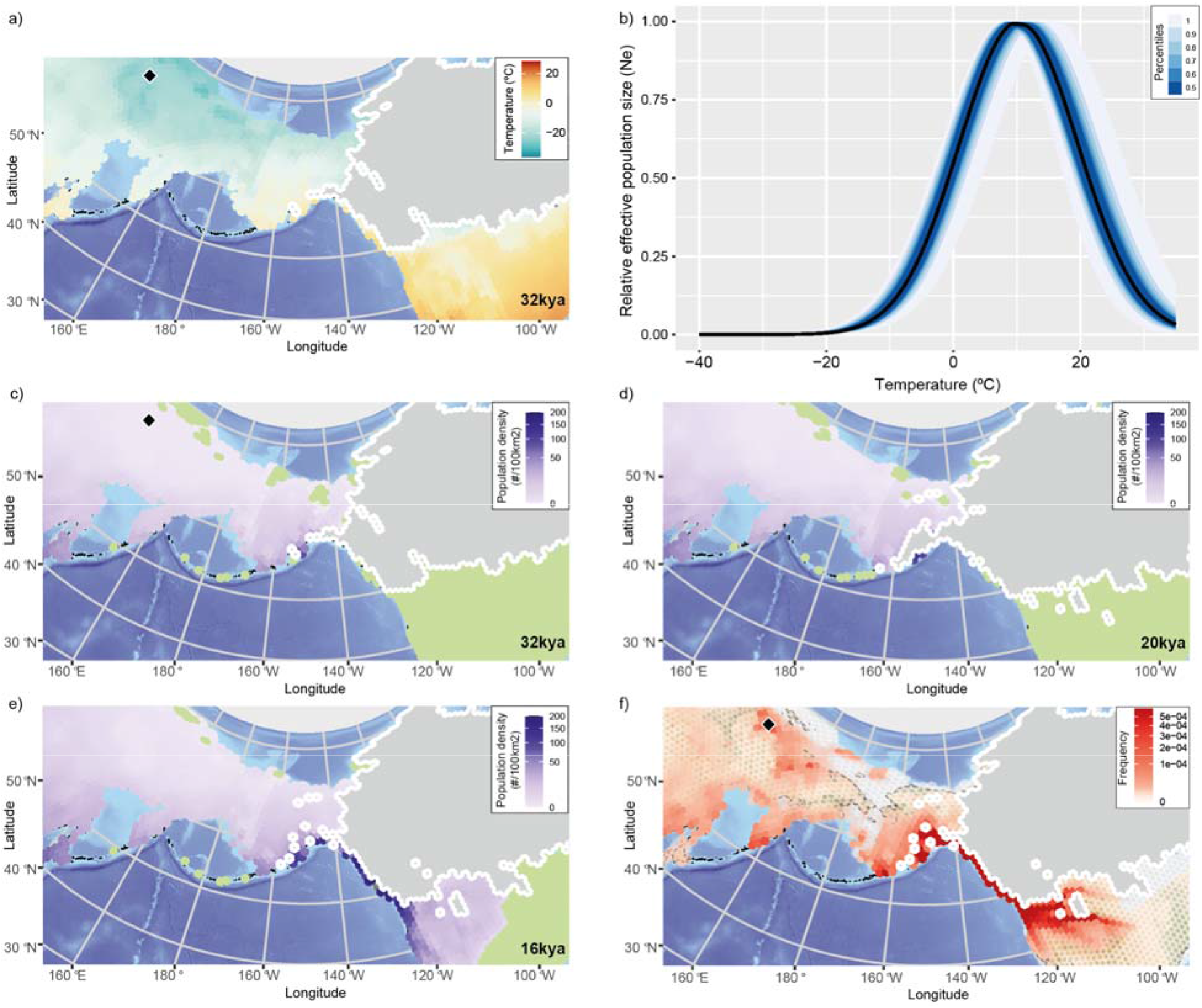
a) Mean annual temperature plot at 32 kya, when the Yana individual (black dot) lived. b) Effect of temperature on the relative population size from the best fitting scenarios, indicating the ability to persist at very low temperatures. The solid black line represents the median of all best fits, whilst the blue shading shows the uncertainty as percentiles. Effective population density across the best fitting scenarios at c) 32 kya, d) 20 kya and e) 16 kya years ago. Ice sheets are shown in gray, uncolonised areas in green. f) Distribution of common ancestor events between 50 and 16 kya: the role of Beringia and the strong founder events during the rapid colonisation along the coastal corridors are shown by a concentration of CA events. Sampled population (Yana) is shown with the same colour used in Fig. 1.

In our model, the colonisation of the Americas requires the coastal corridor along the Pacific coast west of the Cordilleran Ice Sheet^18^ (**Fig. 4e**). Considering the high tolerance to low temperature, the entrance in the Americas started as soon as the corridor was available between 17 and 16kya (**Fig. 5a**), which is compatible with the genetic separation among early native American genomes. We only have high coverage ancient genomes for the western part of North America, but we can explore the predicted gene flow by simulating an additional population in the east (specifically, we chose the ancient Southwestern Ontario sample, ASO^20^). Our model supported the existence of two main streams of gene flow into North America^20^: Ancient-A, including Anzick-1, descending eastwards of the Rocky Mountains; and Ancient B, moving towards the East Coast, reaching ASO (**Fig. 5b**). As we only have one high coverage ancient sample in South America, we are not able to reconstruct the details of the possible admixture of these two streams in that region. However, we identified an initial entrance of South America around 14.6 kya (**Fig. 5c**) with the continent fully colonised by 13.2 kya (**Fig. 5d, Extended Data Fig. 5**). For such a fast colonisation process, the model selected high values for the parameter underlying movement into uncolonised regions (directed mobility, m_d_) combined with an increase in migration speed over time (Δ_speed_) to generate a rapid expansion rate (**Fig. 5e, Extended Data Fig. 6b,f**). We estimated an expansion speed of 4.1 km/year by calculating the shortest path from the opening of the Cordilleran Ice Sheet to Sumidouro5 (Brazil) that had to be crossed in 3.2 kyrs considering the earliest colonist reaching Brazil in our model by 13.8 kya. This date is compatible with some of the archaeological evidence for the earliest human occupation of the continent ^21,22^. Such a speed is within the range of the average distance per year covered by contemporary HG populations^23^.

**Fig. 5:**
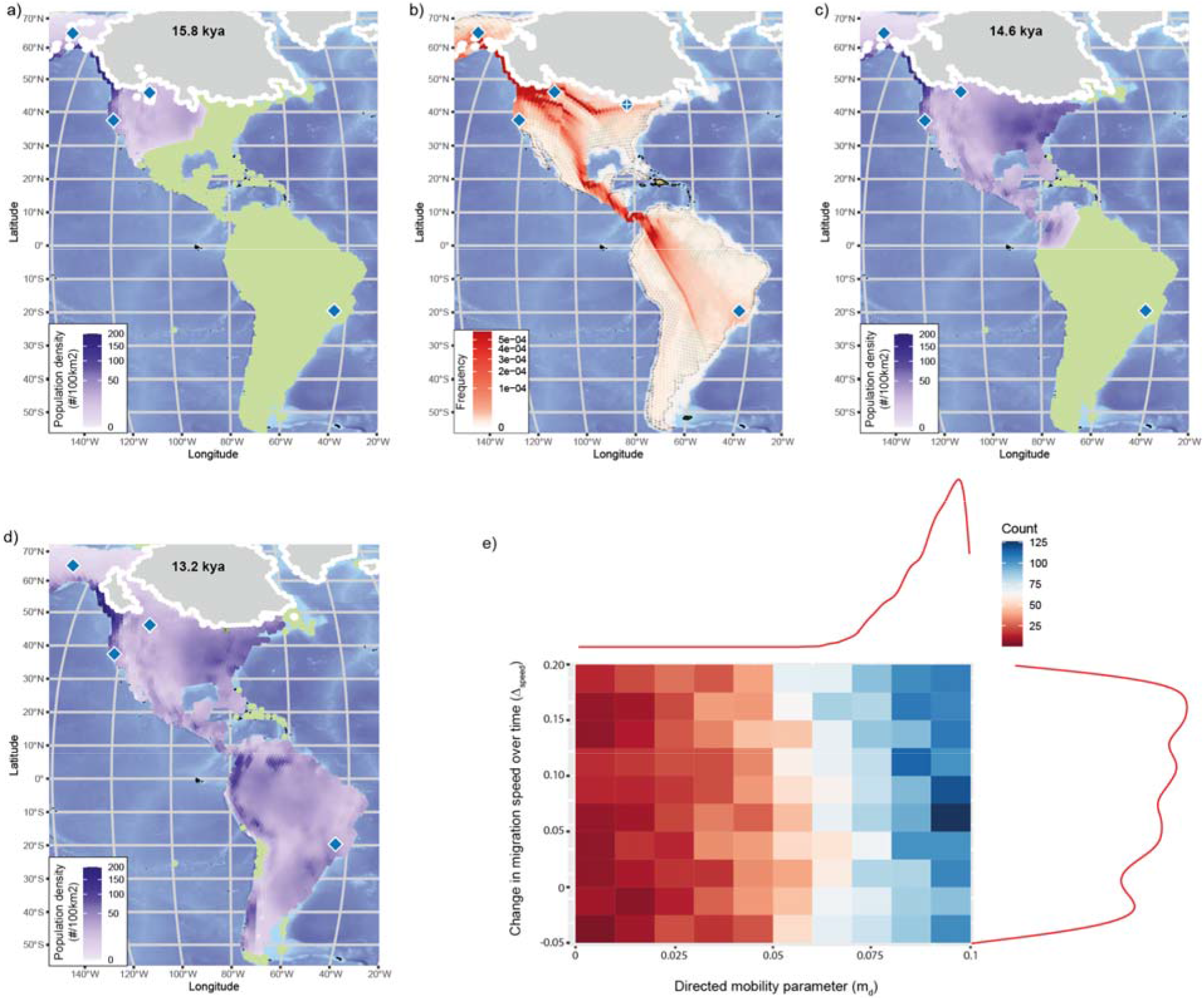
a) Effective population density at the beginning of the colonisation of the Americas (15.8kya) after the opening of the coastal corridor along the Pacific coast west of the Cordilleran Ice Sheet. b) Distribution of common ancestor events for the last 25 kyrs: the two main streams of gene flow in North America represent ANC-A and ANC-B as in^20^. The artificial population representing ancient Southwestern Ontario (ASO) is highlighted with a white cross. c) Effective population density map showing the entrance in South America (14.6 kya) and d) when the continent is fully colonised by 13.2kya. e) The combined effect of change of migration speed over time and directed mobility parameters suggesting high values to generate a rapid expansion rate required to fully colonise South America by 13.2kya. (from the best 0.2% scenarios retained during parameter estimation). The posterior distribution for each parameter is shown on their corresponding axis.

## Discussion

Our model assumes that the link between climate and the demography of HGs did not change significantly following the Out-of-Africa expansion, only allowing a change in migration speed. Given the ability of the model to faithfully reproduce the observed genetic patterns, this suggests that the demography of HGs did not change significantly over that time period. However, whilst the signals that these processes leave in the genomes are strong enough that we were able to retrieve clear peaks in our posteriors (and thus quantify the relative importance of these processes), but the posteriors are still relatively broad (**Extended Data Fig. 6**), implying that the quantities that we used to describe the genomes cannot pinpoint exact values. Importantly, the quantities that we used to describe the differentiation among lineages (*π*, the number of pairwise differences between genomes) are robust to rapid recent expansions, and thus would be blind to changes that might have occurred over the last few thousand years (as one might expect as a result of the interaction between HGs and food producers). In summary, whilst mostly time-invariant demographic rules (except an increase in migration rates) are sufficient to explain the differentiation among ancestral human lineages, this does not imply that the cultural changes that happened during the last 70k years did not have any effect on the demography of different worldwide populations.

In our modelling, we did not attempt to look at different African lineages, as we lack a comprehensive catalogue of past African genetic diversity, and aDNA preservation in Africa is limited to the recent past^24^. We would expect that the factors identified in this study will also play an important role in Africa. However, the deep separation of the Koi San and Mbuti from other African populations^25^ suggests that climate and mountains alone might not be sufficient to explain the spatial structuring in Africa, and that additional processes will have to be included to explain the divergence among African ancestral lineages.

Recent work on aDNA has emphasised how modern Out-of-Africa populations derive from a mixture of distinct ancestral lineages^1^, reflecting high levels of mobility during the Holocene following the advent of agriculture and animal husbandry^2,3^. Here, by formally combining high quality ancient and modern genomes with comprehensive paleoclimate reconstructions, we have shown that those ancestral lineages emerged following the Out-of-Africa expansion as a response to climatic changes and terrain that influenced the demography and mobility of hunter gatherers. Whilst the advent of food production led to large scale movements that we have not investigated here, recent work on migrations during the Neolithic and Bronze Age has shown that climate continued to play an important role during those later periods ^4,26^.

## Methods

### Samples

The dataset published in Maisano Delser et al.^27^ was subset and the full list of included samples is shown in Supplementary Table 1 and 2. From the initial set of 35 samples, three samples (NE1, NE5, ZVEJ31 and LBK) were discarded because they do not represent ancient hunter-gatherer populations. Preliminary analyses were based on 31 samples worldwide distributed (HG_EXT dataset). For spatial analysis, we favoured ancient over modern individuals where both of them were available for the same geographical area (7 modern samples discarded). We also added Bon002 because it represents a proxy for pre-pottery Neolithic in Turkey^28^. The final dataset (HG dataset) for the spatial analysis includes 25 samples (16 ancient and 9 modern individuals) representing hunter-gatherer populations distributed worldwide.

### Genetic summary statistics

#### Neutral loci

We selected 9178 loci of 10kb in length along the genome to represent neutral genetic variation. We first applied to the human genome filters to exclude regions that may be under selection or that could be problematic in terms of assembly quality: more specifically, we filtered out coding regions, conserved elements, recombination hotspots (regions with recombination rates >10cM/Mb), repetitive regions, and regions with poor mapping or sequencing quality. The filters are the same as used in Kuhlwilm et al. (2016)^29^ and were kindly provided by Ilan Gronau. Contiguous intervals of 10kb were chosen on these remaining sites using a sliding window approach. Windows were retained if 7,500 sites or more were present in a panel of 7 modern samples distributed worldwide (see Supplementary Information), and a subset of these were selected with a minimum inter-locus distance of 50kbp. This minimum 50kbp distance between loci was chosen so that the chance of recombination was sufficiently high that loci could be treated as unlinked. Finally, 118 regions were discarded after filtering for coverage and CpG sites leaving 9060 windows.

#### Genetic diversity

The HG_EXT dataset is a subset from^27^ which includes 84,782,047 sites called for 31 samples (16 modern and 15 ancient individuals, see **Extended Data Table 1 and 2**) without any missing data and triallelic site. Number of pairwise differences was calculated with plink v1.9 (--distance 1-ibs allele-ct flat-missing square --allow-no-sex)^30^ and divided by the total number of sites to obtain estimates of whole-genome pairwise π (π_wg_). However, values of π_wg_ for the non UDG-treated samples are not reliable because of the inflation of transitions due to DNA damage and degradation. Therefore, the number of transitions (ts) and transversions (tv) per sample was calculated with bcftools stats v1.6^31^. The ratio ts/tv was calculated per sample and plotted in R v3.6.3^32^. Transversions only were extracted from HG_EXT dataset into a new vcf file and the number of pairwise differences was recalculated with plink v1.9 (--distance 1-ibs allele-ct flat-missing square --allow-no-sex)^30^p. The average ts/tv was calculated across all samples but non UDG-treated samples (7 in total: Bichon, WC1, Yana1, Kolyma1, Sumidouro5, Anzick-1 and Mota) and Loschbour as it shows an unusual low ts/tv ratio. Estimates of pairwise π for the non UDG-treated samples and Loschbour were calculated by rescaling the number of pairwise differences obtained from the transversions only (diff_tv) to π_wg_ using the mean ts/tv ratio (see above) with the formula 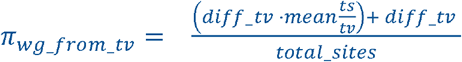. We also assessed the correlation between π_wg_ and π_wg_from_tv_ generated with the formula above for modern and ancient UDG-treated samples (see Supplementary Information).

We subset the HG_EXT for the neutral loci identified above using tabix v1.9^31^ and we retained 9,618,572 sites. At this stage, we called these sites as haploid in Bon002 with pileupCaller^33^ because the coverage did not allow for calling diploid genotypes reliably. When merging Bon002, 494,784 sites were discarded because of missing data bringing the total number of sites to 9,123,788. These sites have been split in the 9069 windows identified previously averaging ∼1kb per window. We converted the diploid vcf file into a haploid vcf file to match the approach used in msprime^34,35^ where haploid chromosomes are simulated (convert_diploid_into_haploid_vcf_gz.sh, available in the GitHub repository). We then rechecked the number of transitions and transversions per sample and the ts/tv ratio was recalculated based on the same samples set used for whole-genome data. Estimates of π_wg_ for ancient non UDG-treated samples and Loschbour were rescaled from values based on transversions only (π_wg_from_tv_) with the same approach described above. The correlation between π_windows_ and π_wg_was also calculated (see Supplementary Information).

#### Spatial Simulations

Spatial simulations were performed in CISGeM which stands for Climate Informed Spatial Genetic Models. Within this framework, information from climatic reconstructions is used to drive the local demography within a spatially explicit metapopulation model, which in turn is used to simulate genetic data (**Extended data Fig. 7**).

#### Carrying capacity and demographic model

CISGeM’s demographic module consists of a spatial model that simulates long-term and global growth and migration dynamics of anatomically modern humans (AMHs). These processes depend on a number of parameters (see Supplementary Table 3), which we later estimate statistically based on empirical genetic data.

The model operates on a global equal area hexagonal grid of 40,962 cells that represent the whole world (the distance between the centers of two hexagonal cells is 120.6 ±7.6 km; the variation is due to the earth having a spheroid rather than a perfect spherical shape and the grid not being perfectly regular). Each model time step represents 25 years, appropriate as the generation time in AMHs. Each time step of a simulation begins with the computation of the carrying capacity of each grid cell, i.e. the maximum number of individuals theoretically able to live in the cell for the environmental conditions at the given point in time. Specifically, we consider the impact of precipitation and temperature on population density. Precipitation and temperature are based on global climate model (HadCM3) simulations of the last 120ka^36^ that have been extended further back in time using a linear regression approach^37^. The data set has been bias-corrected^38^ and is available at a spatial resolution of 0.5°x0.5°. The climate data represent the climatological mean for every 1ka throughout the last 300ka. It has been spatially interpolated from the regular 0.5° lon/lat grid to the hexagonal grid, and temporally interpolated onto 25yr time steps (linearly) using the climate data operators^39^. We require precipitation to be above a minimum threshold (*P*_*crit*_, below which humans cannot survive), with population density increasing with increasing precipitation above that level. For temperature, we model human tolerance by considering an optimal annual mean temperature (*T*_*opt*_, at which the highest population density can be achieved) and a scaling parameter (*T*_*tol*_) which defines how well humans can cope with deviations from *T*_*opt*_. The larger *T*_*tol*_, the more tolerant humans are, thus leading to relatively large population densities despite suboptimal temperatures. The carrying capacity (in effective individuals, Ne) in a grid cell *x* at a time, *t* was modelled as:

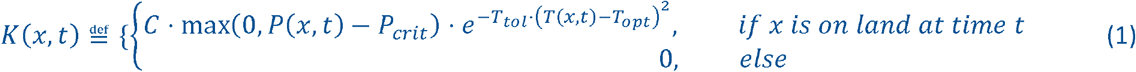

Where *P* (*x, t*) and *T* (*x, t*) denote the annual precipitation (in mm year^-1^) and the mean annual temperature (in °C) in the cell *x* at time *t*, respectively. is a scaling constant. We confirmed that this relationship is biologically plausible by verifying that it provides a good fit to census population sizes for modern hunter-gatherer groups^40^ (**Extended data Fig. 8**). However, note that the parameters were fitted to the genetic data (see below); the census population sizes of modern hunter gatherers were only used to test that the shape of the relationship was realistic.

The estimated carrying capacities are used to simulate spatial population dynamics as follows. We begin a simulation by initialising a population in a grid cell *x*_0_ at a point in time *t*_0_ with K (*x*_0,_ *t*_0_) individuals. For our simulations we chose *x*_0_ to be in East Africa (we arbitrarily chose the cell closest to 26.5° E, 9.7° N as in ^9^). The exact location has no impact on the simulations, as we were not concerned with the within-Africa population structure and we only have one African genome to represent the split between African and Out-of-Africa lineages.

At each subsequent time step between *t*_o_ and the present, CISGeM simulates two processes: the local growth of populations within grid cells, and the spatial migration of individuals across cells. Similar to previous work^7,41,42^, we used the logistic function to model local population growth in humans, estimating the net number of individuals by which the population of size *N* (*x, t*) in the a *x* at time *t* increases within the time step as

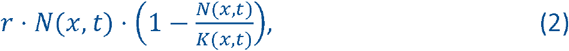

where *r* denotes the intrinsic growth rate. Thus, growth is approximately exponential at low population sizes, before decelerating, and eventually levelling off at the local carrying capacity. Numerically, the ratio of the above term and some positive whole number *n_steps* was applied *n_steps* consecutive times to the relevant population size, where *n_steps* was chosen such that *r/n_steps* is small enough to ensure the stability of the logistic map for the range of growth rates *r* considered in our simulations.

Across-cell migration is modelled as two separate processes, representing a non-directed, spatially uniform movement into all neighbouring grid cells on the one hand, and a directed movement along a resource availability gradient on the other hand. For both of them, movement between two grid cells is reduced when it involves crossing mountains. Under the first movement type, the number of individuals migrating from a cell *x*_1_ into a neighbouring cell *x*_2_ is estimated as

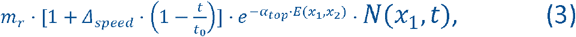

where *α*_*top*_,, speed, *t*_0_, *m*_*r*_ are parameters, and where *E* (*x*_1,_ *x*_2_) is a measure of the altitude that needs to be covered between cells *x*_1_ and, which we defined as follows. We used a very-high-resolution (1 arc-minute) global elevation and bathymetry map (ETOPO1^43^) and determined, for each pair of neighbouring cells *x*_1_ and *x*_2_, the altitude profile along the straight line between the geographic cells. We then defined *E* (*x*_1,_ *x*_2_) as the sum of the absolute values of all altitude changes along the line. This assumes that descends have the same effect in terms of reducing movement rates as ascends; in particular we have *E* (*x*_1,_ *x*_2_) = *E* (*x*_1,_ *x*_2_). Δ*speed* represents the change of migration speed over time, and is scaled by the duration of the simulation 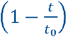 at time *t* (note that *t* is a negative quantity as it represents the number of generations before present). This mechanism in Eq. (3) is equivalent to a spatially uniform diffusion process, which has previously been used to model random movement in AMHs^7,41^. Under the second movement type, an additional number of individuals moving from a grid cell *x*_1_ to a neighbouring cell *x*_2_ is estimated as

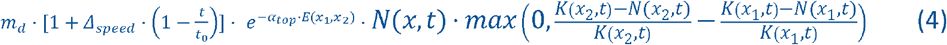

The number 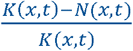 represents the relative availability of unused resources in the cell *x* at time *t*, equalling 1 if all natural resources in *x* are potentially available for humans (*N* (*x, t*) = 0), and 0 if all resources are used (*N* (*x, t*) = *K* (*x, t*)). Thus, individuals migrate in the direction of increasing relative resource availability, and the number of migrants is proportional to the steepness of the gradient. The distinction between directed and non-directed movement allows us to examine to which extent migration patterns can be explained by random motion alone or requires us to account for more complex responses to available resources. The coefficient *m*_*d*_ is a parameter.

For some values of the mobility parameters *m*_*r*_ and *m*_*d*_, it is possible for the calculated number of migrants from a cell to exceed the number of individuals in that cell. In this scenario, the number of migrants into neighbouring cells are rescaled proportionally such that the total number of migrants from the cell is equal to the number of individuals present.

Similarly, it is in principle possible that the number of individuals present in a cell after all migrations are accounted for (i.e. the sum of local non-migrating individuals, minus outgoing migrants, plus incoming migrants from neighbouring cells) exceeds the local carrying capacity. In this case, incoming migrants are rescaled proportionally so that the final number of individuals in the cell is equal to the local carrying capacity. In other words, some incoming migrants perish before establishing themselves in the destination cell, and these unsuccessful migrants are not included in the model’s output of migration fluxes between grid cells. In contrast, non-migrating local residents remain unaffected in this step. They are assumed to benefit from a residential advantage^44^, and capable of outcompeting incoming migrants.

CISGeM’s demographic module outputs the number of individuals in each grid cell, and the number of migrants between neighbouring grid cells, across all time steps of a simulation. These quantities are used to reconstruct genetic lineages.

Once a demography has been generated, gene trees are then simulated (the genetic code borrows heavily from msprime^34,35^). This process depends on the population dynamics recorded during the demography stage and assumes local random mating according to the Wright-Fisher dynamic. From the present, ancestral lineages of sampled individuals are traced back through generations, recording which cell each lineage belongs to. At every generation, the lineages are randomly assigned to a gamete from the individuals within its present cell. If the assigned individual is a migrant or coloniser, the lineage is moved to the cell of origin for that individual. Common ancestor events happen when two lineages are assigned to the same parental gamete and they are then merged into a common ancestor lineage. This process is repeated until all the lineages have met. If multiple lineages are still present at the time when the demography was initialised, the remaining lineages enter a single ancestral population (with fixed population size K_0_), and a coalescent model is used to estimate the timing of additional common ancestors events to close the tree (see^34^ for an example of using hybrid models where the coalescent is used to close trees generated by an initial Wright-Fisher dynamics).

To match our data design, we simulated 9060 non-recombining loci of 1kb each. For each locus, mutations were added to the gene tree with a mutation rate *µ* = 1 · 10^−8^ site/generation^6^.

Parameter space was explored with a Monte Carlo sweep. Each parameter was randomly sampled from a uniform prior distribution (see Supplementary Table 4 for ranges). We generated a total of 3,129,979 simulations, out of which 252,860 reached all our sample locations at the required dates as estimated for the fossil remains.

#### Parameter estimation

CISGeM output files generated from the Monte Carlo sweep were processed with rcisgem v1.0, an R package developed to create, edit and process files for CISGeM. Individuals populations were aggregated into ancestral lineages: North-Central Asians (“nca” includes Yana1, Xibo, Mansi and Uts_Ishim), North-Eastern Asians (“nea” includes Kolyma1, Oroqen, Ulchi and Yakut), South Asia (“sa” includes Irula), South-East Asians (“sea” includes Papuan, Jehai and Australian), Western Hunter Gatherers (“whg” includes Sunghir III, Bichon, SF12, Loschbour and ZVEJ25), Near East (proto-Neolithic Bon002), Caucasus Hunter Gatherers (“chg” includes KK1 and Iranian early Neolithic WC1), Africa (“af” includes Mota) and Native Americans (“am” includes Anzick-1, USR1, Sumidouro5 and AHUR_2064). Pairwise π comparisons between these lineages for both observed and simulated data were calculated with the function compute_abc_sumstats.R from the package rcisgem. Approximate Bayesian Computation based on regression is limited to a small number of statistics (usually not more than ten^45^). We therefore focussed on a number of comparisons among adjacent lineages that capture the full structure of human genetic diversity: Africa vs North-Central Asia (af_nca), Western Hunter Gatherers vs North-Central Asia (whg_nca), Americas vs North-Eastern Asia (am_nea), Europe vs Caucasus Hunter Gatherers (whg_chg), North-Central Asia vs South Asia (nca_sa), Africa vs South-East Asia (af_sea), Africa vs North-Eastern Asia (af_nea), Africa vs Western Hunter Gatherers (af_whg) and Near East vs Western Hunter Gatherers (ne_whg). Parameters estimation was performed in an Approximate Bayesian Computational (ABC) framework with the library abc v2.1^46^. The best 2% of simulations (i.e. tolerance=0.02), based on the sum of Euclidean distances between observed and simulated summary statistics, were used for the ABC analysis, using a local linear regression to estimate the posterior distributions. Median and mode estimates with 95% credible intervals of the posteriors are reported in **Extended Data Table 4**. Posterior distributions (**Extended Data Fig. 6**) were plotted with the function plot_posterior.R from rcisgem. Simulations retained by the ABC approach were also highlighted in the plots showing the pairwise comparison across all our summary statistics for all simulations (blue dots in **Extended Data Fig. 5**). Histograms of arrival times per population were plotted with the function plot_arrival_times.R from rcisgem (**Extended Data Fig. 5**).

#### Model fitting

We then assessed whether the model was able to recreate the observed genetic diversity by plotting pairwise comparison across all our summary statistics (**Extended Data Fig. 5**). An a priori Goodness-of-fit PCA test was also performed (**Extended Data Fig. 2**) using the *gfitpca* function from R the package abc v2.1^46^ to capture and plot the two first components obtained with a principle component analysis of the simulated summary statistics. We used a *cprob* value of 0.2, 0.35 and 0.5 leaving a different proportion of points from the model outside the displayed envelope (so keeping the best 80%, 65% and 50% points within the envelope). The observed summary statistics is then marked to check that it is contained within these envelopes, thus suggesting a good fit of the model. We also used the *gfit* function to confirm that our model outperformed a series of null models. In this function the goodness of fit test statistic, or D-statistic, is the median Euclidean distance between the observed summary statistics and the nearest (accepted, with the same threshold of 0.02 used during the parameter estimation) summary statistics. For comparison, a null distribution of D is then generated from summary statistics of 1000 pseudo-observed datasets. A goodness of fit p-value can then be calculated as the proportion of D based on pseudo-observed data sets that are larger than the empirical value of D. Therefore, a non-significant p-value suggests that the distance between the observed and accepted summary statistics is not larger than the expectation, confirming that the model fits the observed data well. Both analyses have been performed on a random subset of 250,000 simulations.

#### Demographic scenarios and common ancestor events

Demography output files were generated for the 5058 simulations retained by the ABC during the parameter estimation. For each simulation, we extracted the values of effective population size per cell per generation. Weights from the retained simulations during the parameter estimation (5058) were extracted and rescaled between 0 and 1 (*w*=*w*/sum(*w*)). Effective population size values for each simulation were then multiplied by rescaled weights calculated above and the weighted average was then computed as a sum across the 5058 simulations. We then converted the number of individuals into population density per 100 km^2^ dividing by the grid cell area (124.5 × 100 km^2^) using cdo v1.9.6^39^. Effective densities for present day were compared to estimates based on ethnographic censuses^47^ using a correlation corrected for spatial autocorrelation, using the modified.ttest function from the R library SpatialPack.For the simulations retained (5058) during the parameters estimation, we extracted the time and location of common ancestor events and plotted them as density map.

#### Connectivity with Africa for different temperature tolerances

For a range of tolerance values, we tested, every 500 years, whether there was a direct link between Africa and Eurasia. Connectivity was deemed possible when there was a continuous set of neighbouring inhabited demes that connected Africa all the way to the region outside the Arabian Peninsula, without any breaks. Technically, this was achieved by representing the demes as a graph of connected nodes, using Dijkstra’s algorithm to find the shortest path (if it existed) between African and out of Africa nodes.

## Acknowledgements

P.M.D. was supported by funding from the HERA Joint Research Programme “Uses of the Past” (CitiGen), the European Union’s Horizon 2020 research and innovation programme under Grant Agreement 649307. P.M.D., A.M., M.K., R.B., E.F.M., A.E., M.L. and J.H. were supported by ERC Consolidator Grant 647797 ‘LocalAdaptation’. E.R.J. was supported by a Herchel Smith Research Fellowship. M.P. and I.I.R. were supported by a MRC scholarship. V.S. was supported by the Gates Cambridge Trust. E.T. was supported by a Whitten scholarship. Z.X. was supported by a China Scholarship Council and Cambridge Trust.

## Author Contributions

A.M. designed and supervised the project. M.K. and R.B. generated the paleoclimate reconstruction. A.M., M.K. and R.B., with help from A.E. and V.S., wrote the updated version of CISGeM. P.M.D and E.R.J. collected and processed the genetic data with help from A.H., M.P., D.G.B. and L.C.. P.M.D. performed the relevant genetic analyses and spatial modelling, with help of M.V.. A.M., P.M.D., E.R.J., J.H., and E.F.M. developed the R package “rcisgem”. E.T., Z.X. and A.H. developed the tests for both CISGeM (with help from A.M. and P.M.D.) and rcisgem. M.L. collated and verified the information on modern and ancient samples. G.S. set up the continuous integration for CISGeM. E.P. contributed to develop the effect of mountain ranges. I.I.R. contributed to develop tracking the common ancestor events. A.M. and P.M.D. wrote the manuscript with input from all collaborators. E.F.M. helped to generate figures and maps.

## Data Availability

Cisgem, together with its associated R library rcisgem, will be released as open software on Github upon acceptance of this paper. All input files to rerun the analyses, as well as the code to process the outputs, will be provided on the Open Science Framework.

## Extended data figure and table legends

**Extended Data Fig. 1.**
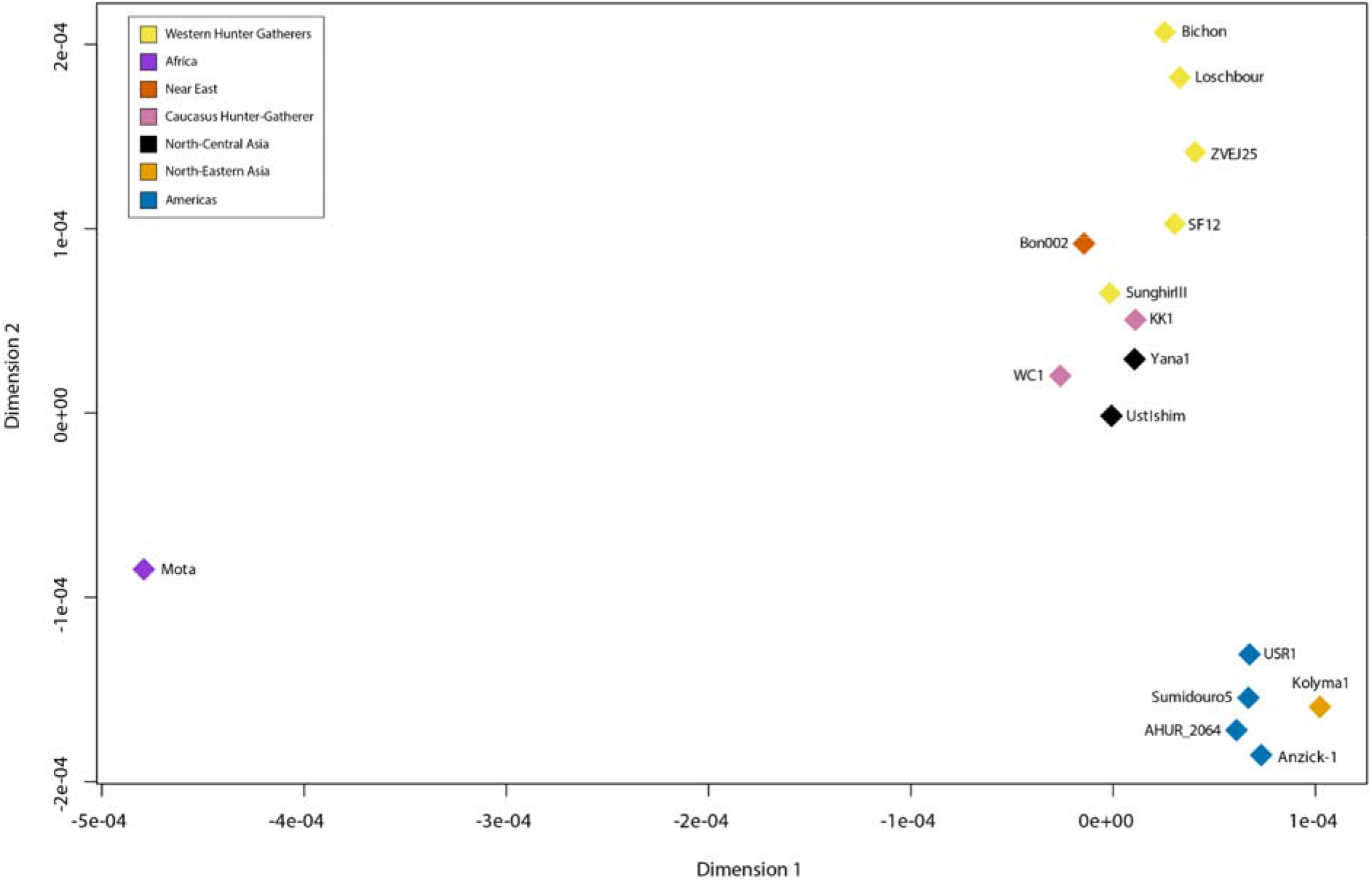
Multidimensional scaling representing the genetic relationships between ancient samples. Whilst Yana1 and Kolyma1 are geographically close, genetically they are distinct: Yana1 shows a greater genetic similarity to Central Asia population while Kolyma1 clusters with North-Eastern Asia and American populations.

**Extended Data Fig. 2.**
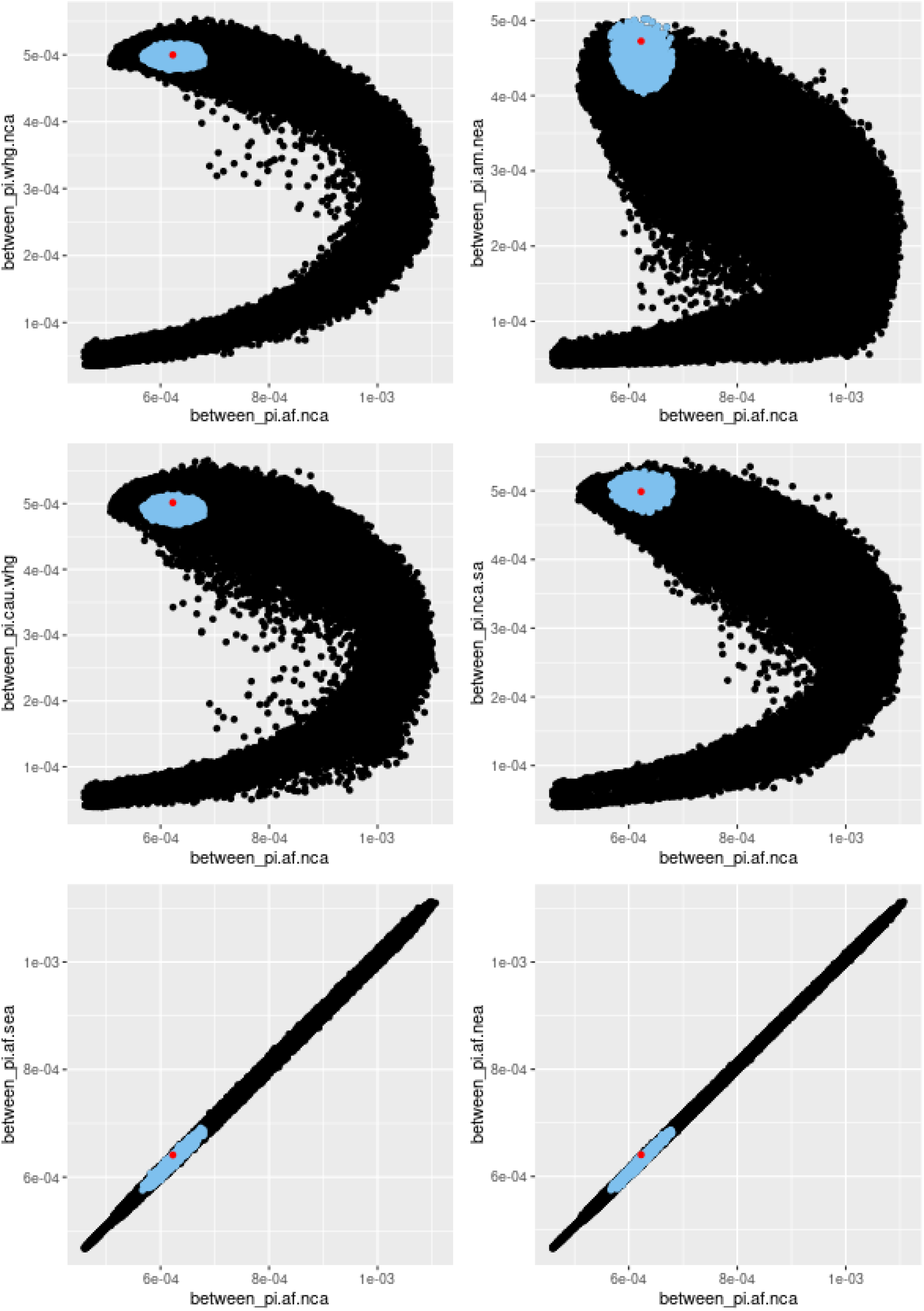
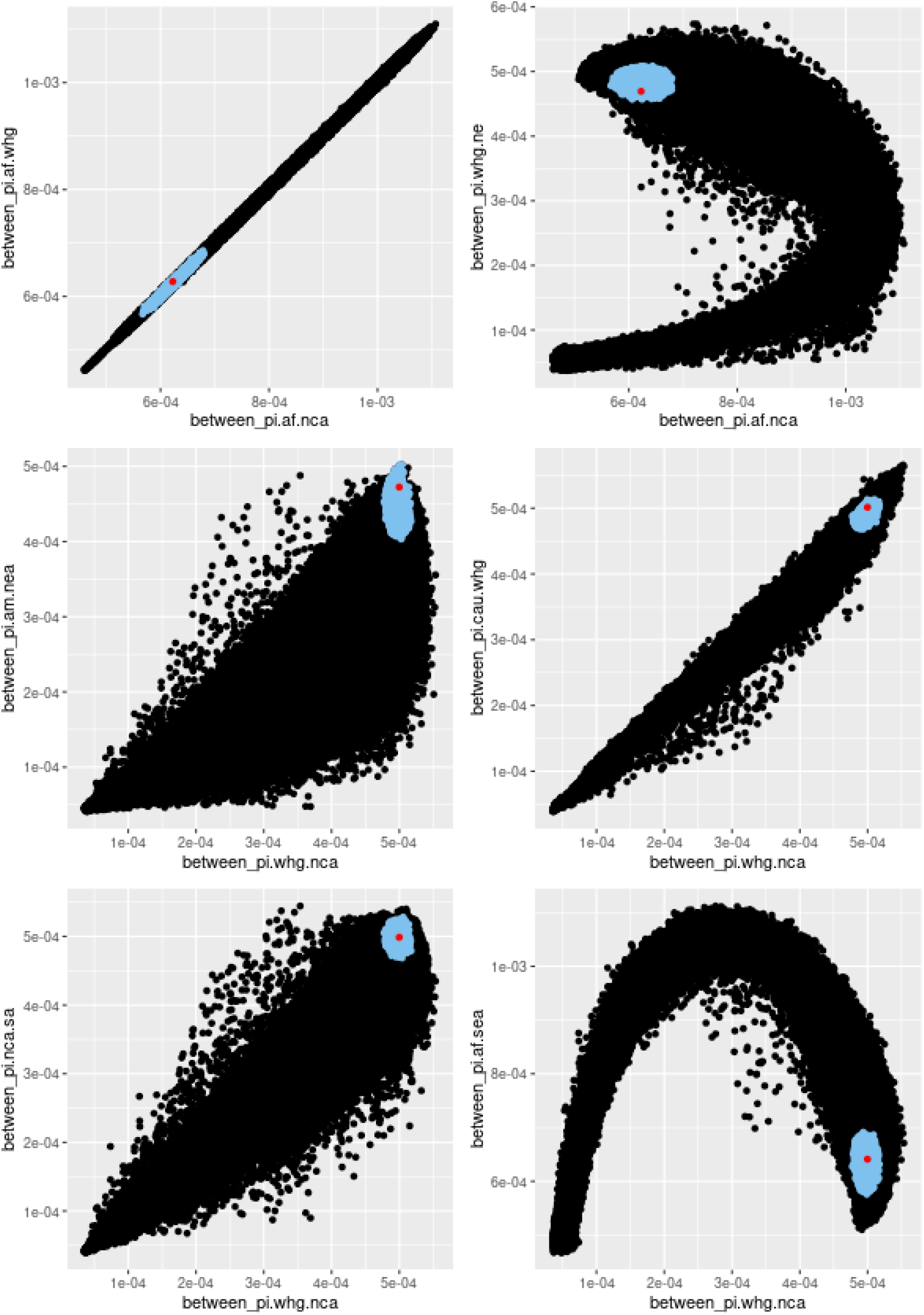
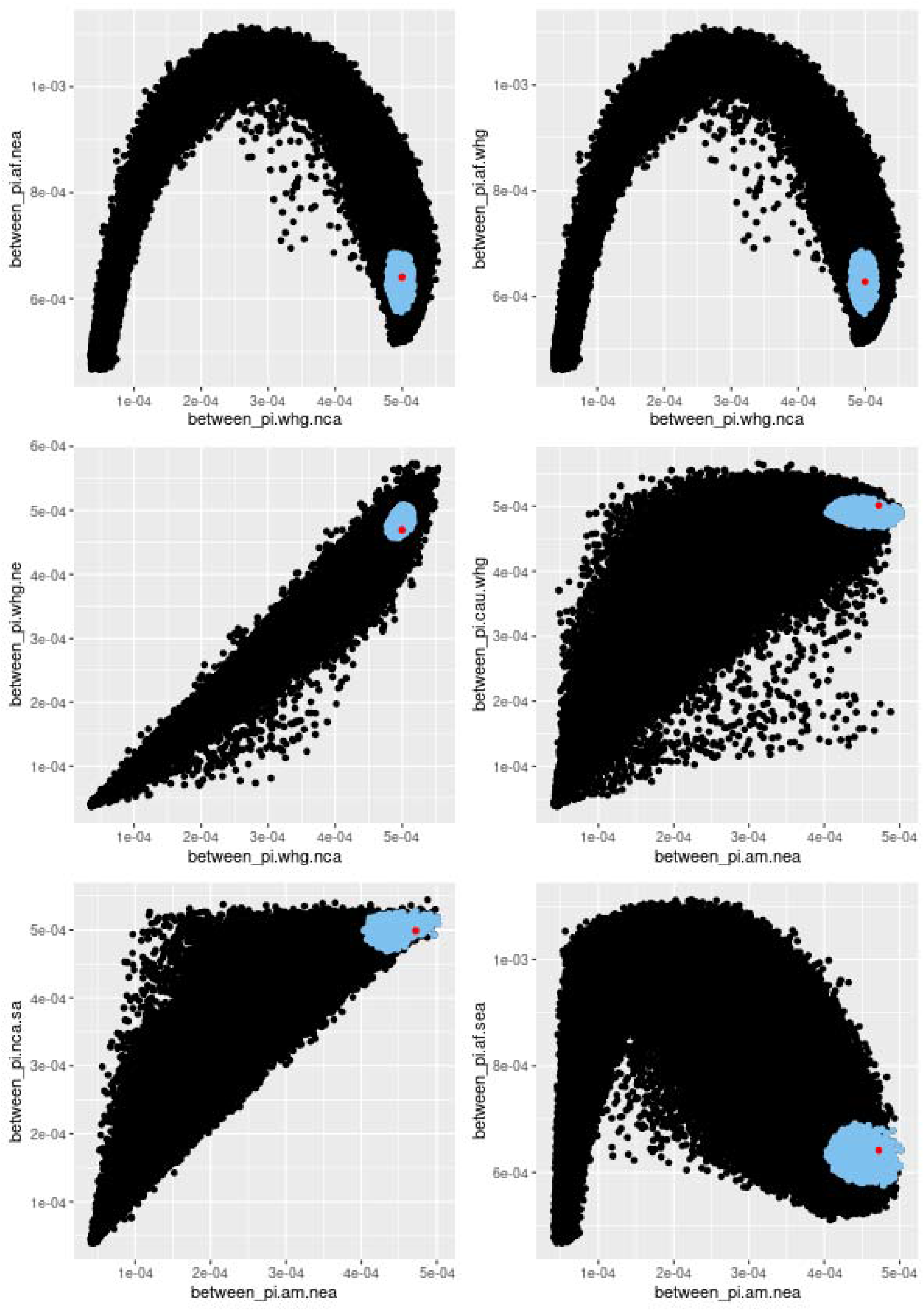
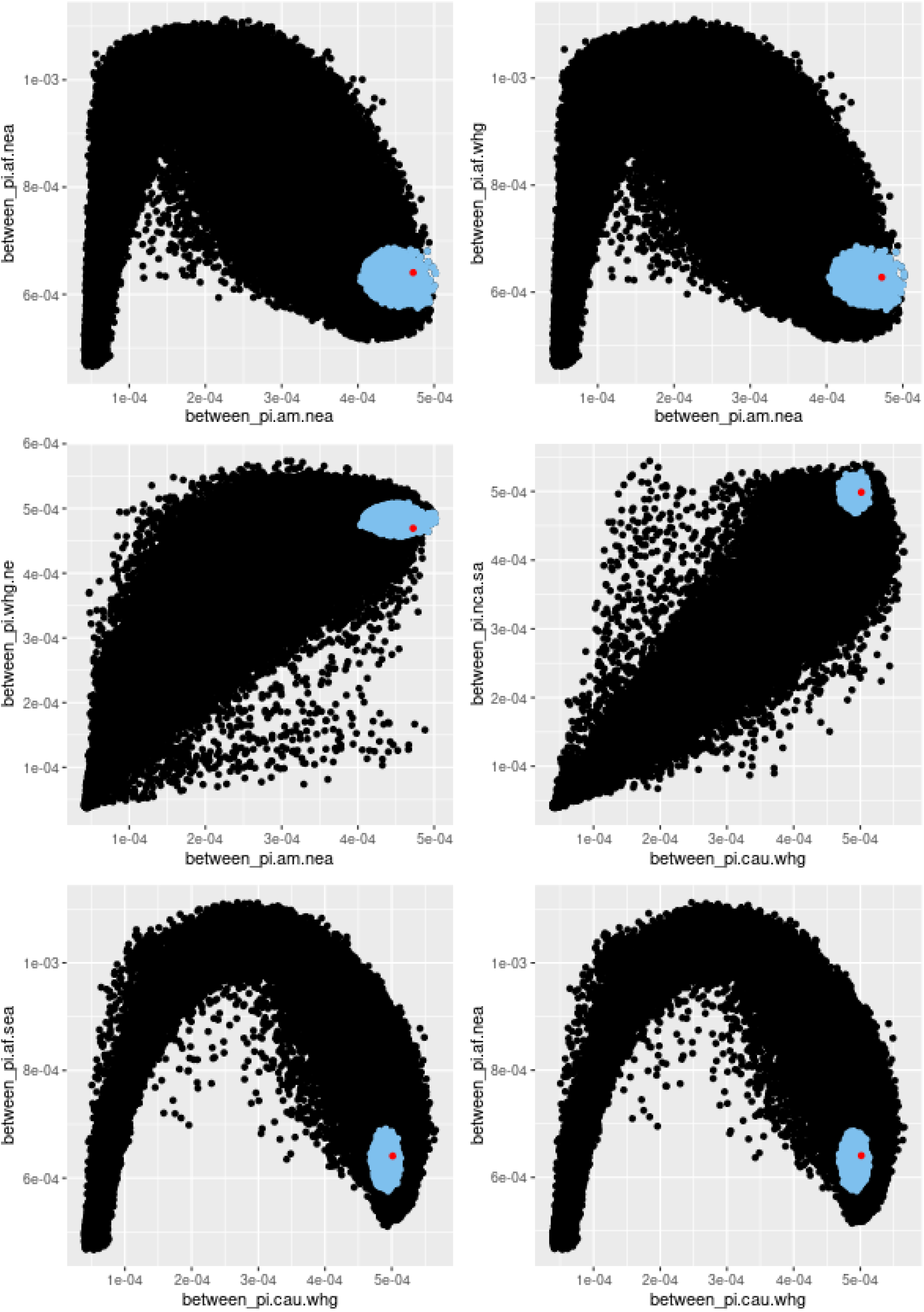
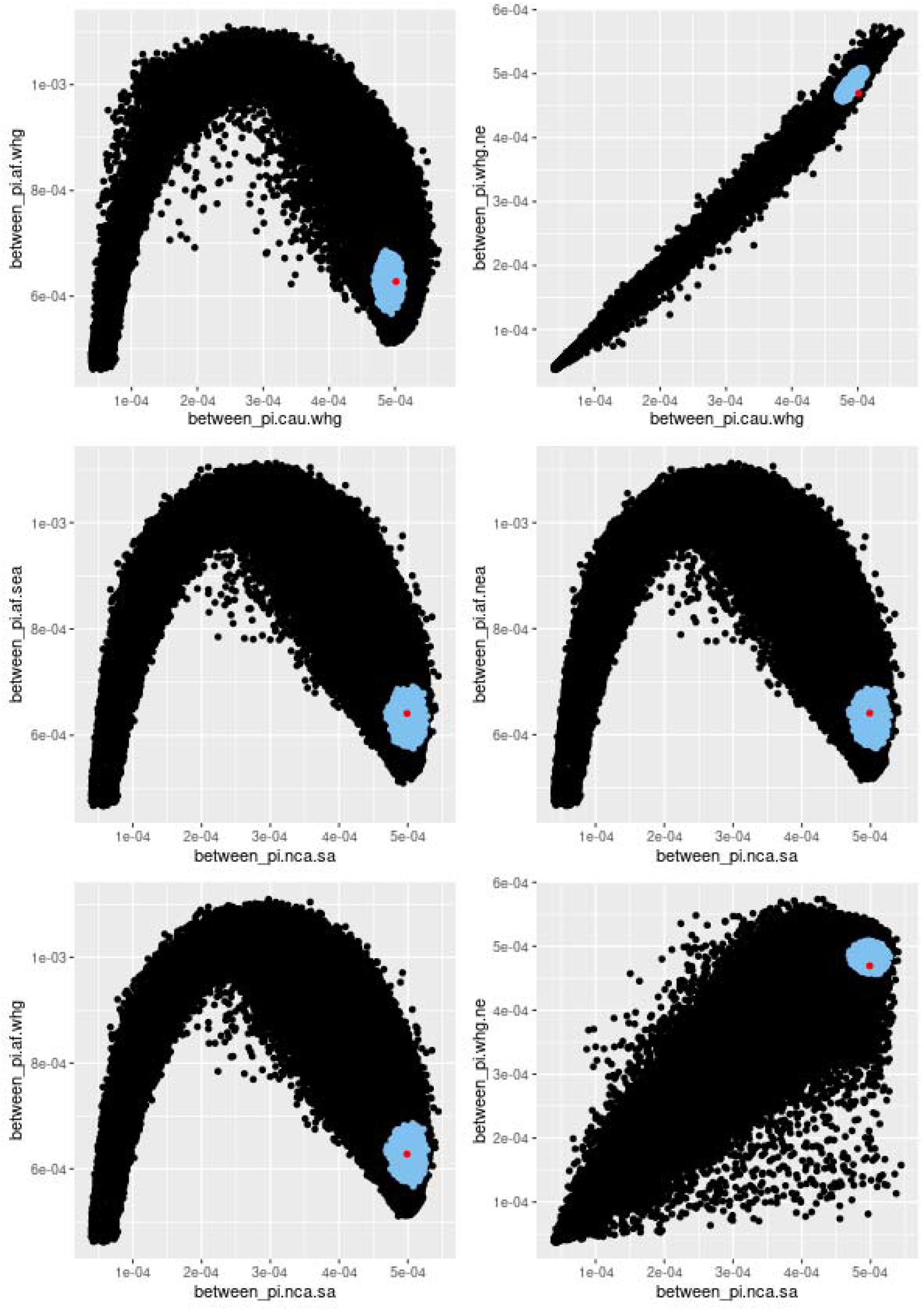
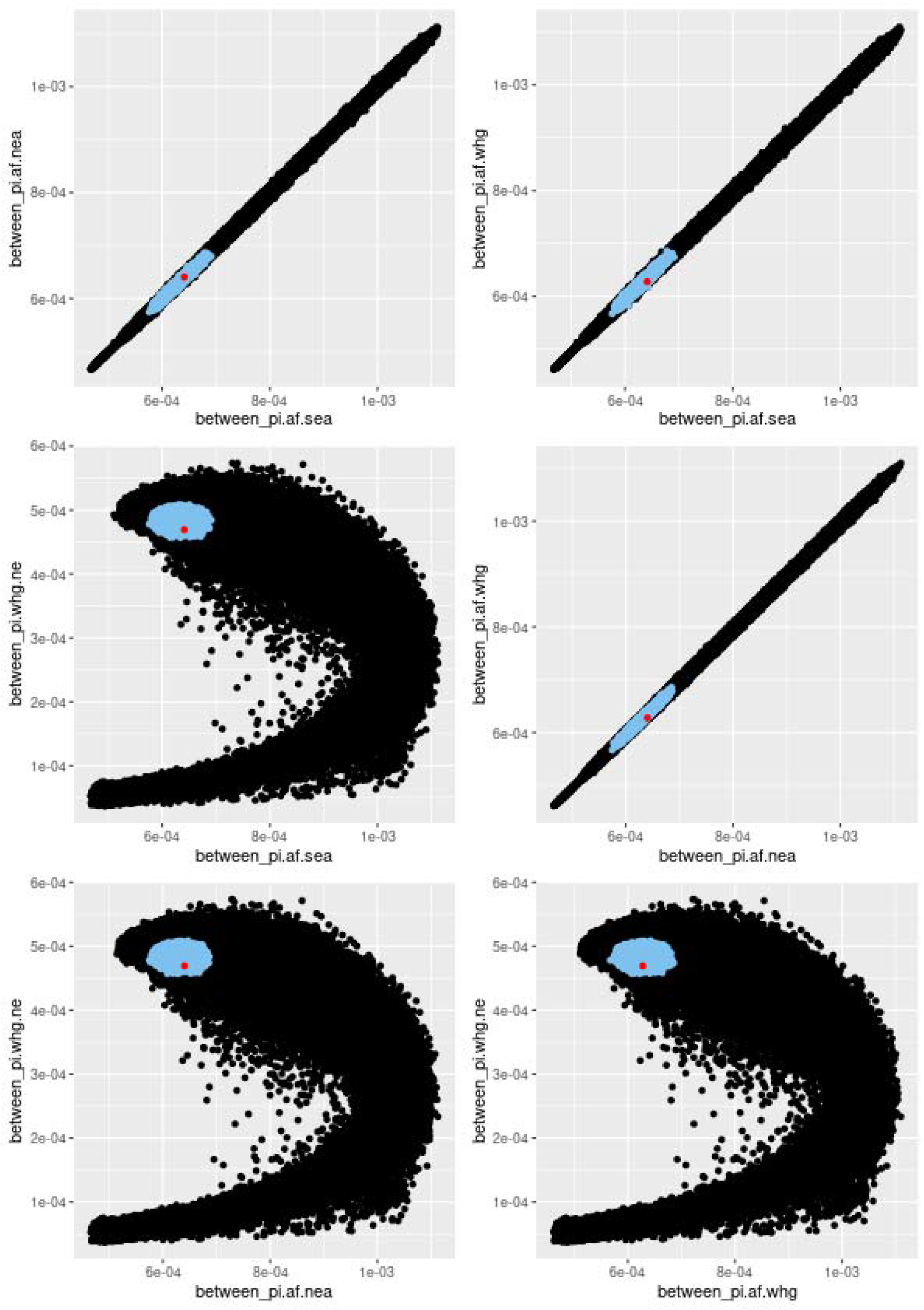
Pairwise comparisons across all summary statistics. For all comparisons, the model is able to recreate the observed genetic diversity as indicated by the fact that the observed values (in red) fall within the simulations (black dots), and also within the subset of best simulations retained during the ABC parameter estimation (in blue), indicating a good fit of the model.

**Extended Data Fig. 3.**
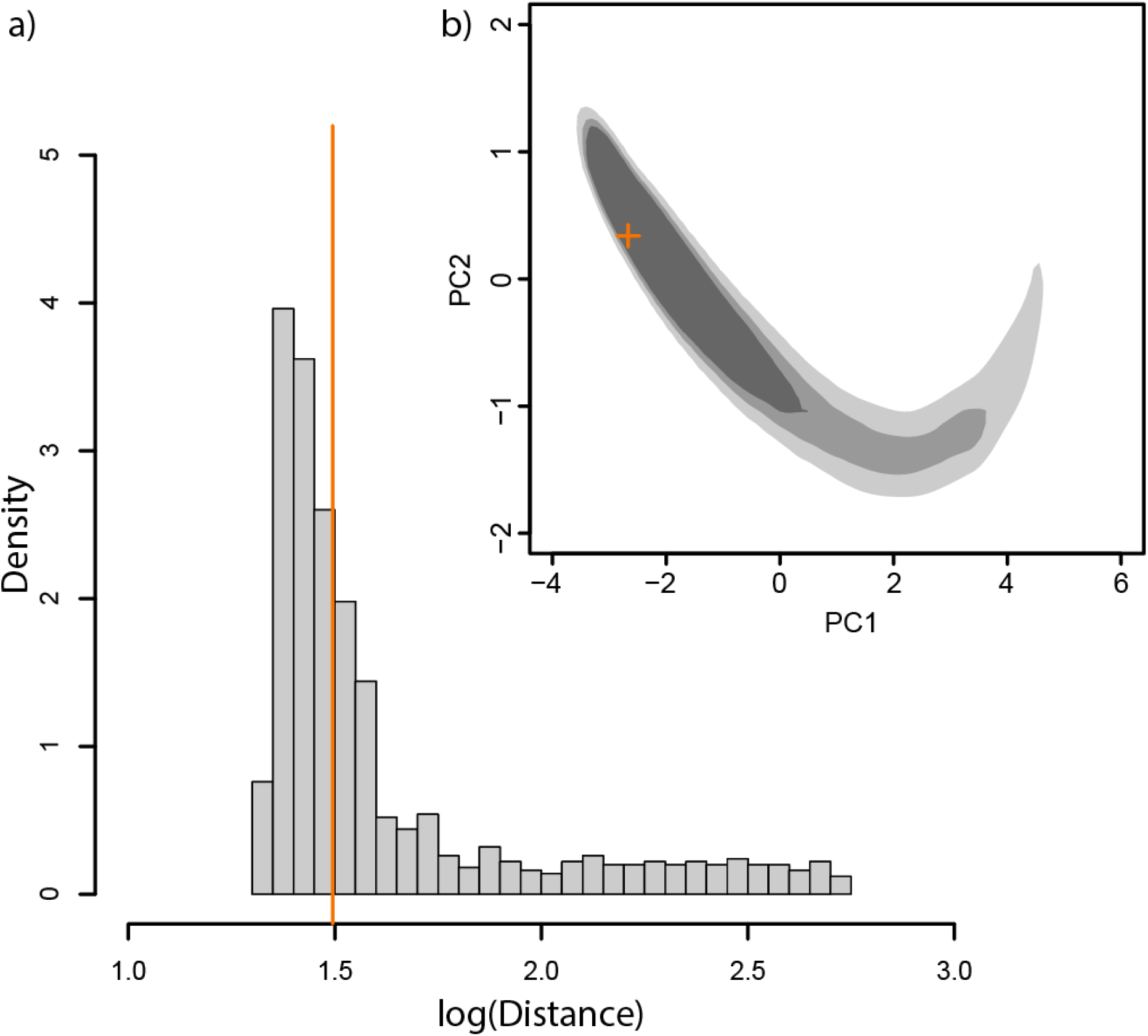
Goodness-of-Fit test. a) histogram representing the null distribution of the logarithmic median Euclidean distance of 1000 pseudo-observed datasets to sets of the closest 2% simulations for each of them. The orange line represents the distance between the observed data and closets 2% simulations (i.e. those retained for ABC parameter estimation with a tol=0.02). A goodness of fit p-value of 0.465 was obtained, indicating that the distance between the observed and retained summary statistics is not larger than the expectation, thus confirming that the model fits the observed data well. b) An a priori Goodness-of-fit PCA test retaining the best 80%, 65% and 50% point in the envelopes from lighter to darker shade respectively. The observed summary statistics (in orange) is contained in all the three envelopes further indicating a good fit of the model.

**Extended Data Fig. 4.**
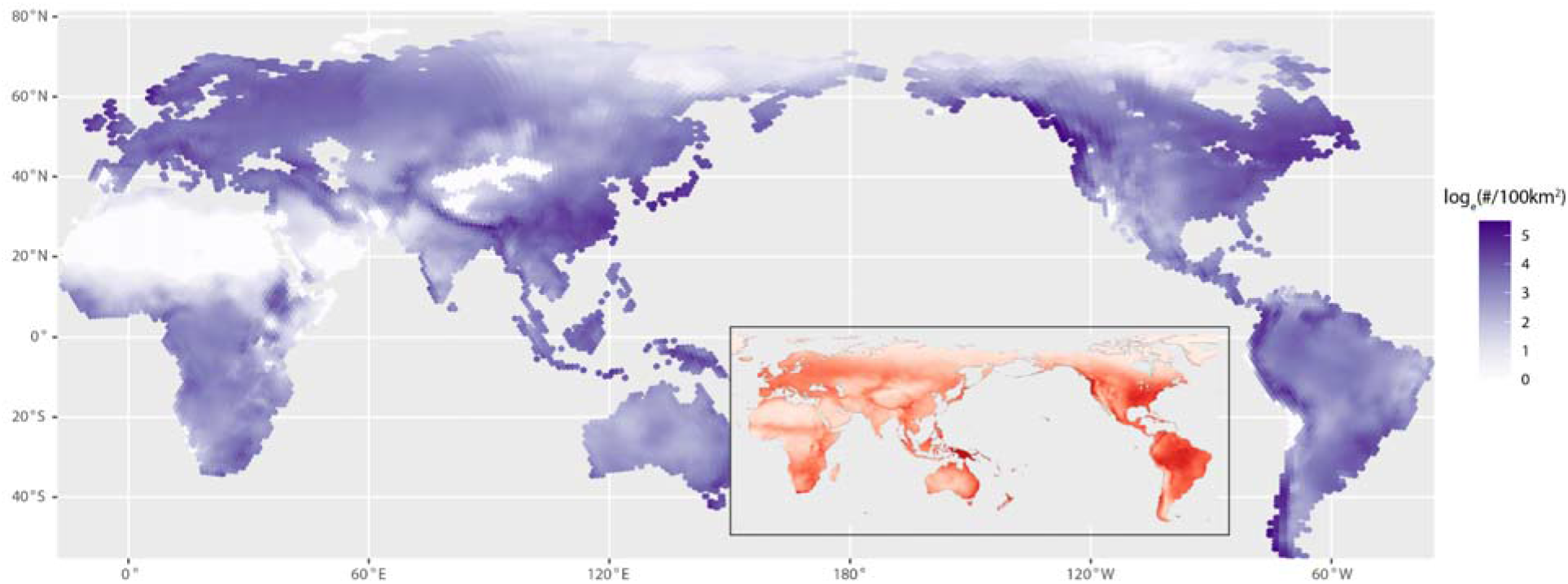
Population density calculated from the weighted mean effective population size (N_e_) of the simulations (5058) retained during the parameter estimation. Inset showing the census sizes from ethnographic censuses (N_census_) from^47^. Despite the differences in data sources and modelling approaches, the two distributions show a high spatial congruence (Pearson’s *r* corrected for spatial autocorrelation: 0.61; p<0.001), with similar regions of high and low densities.

**Extended Data Fig. 5.**
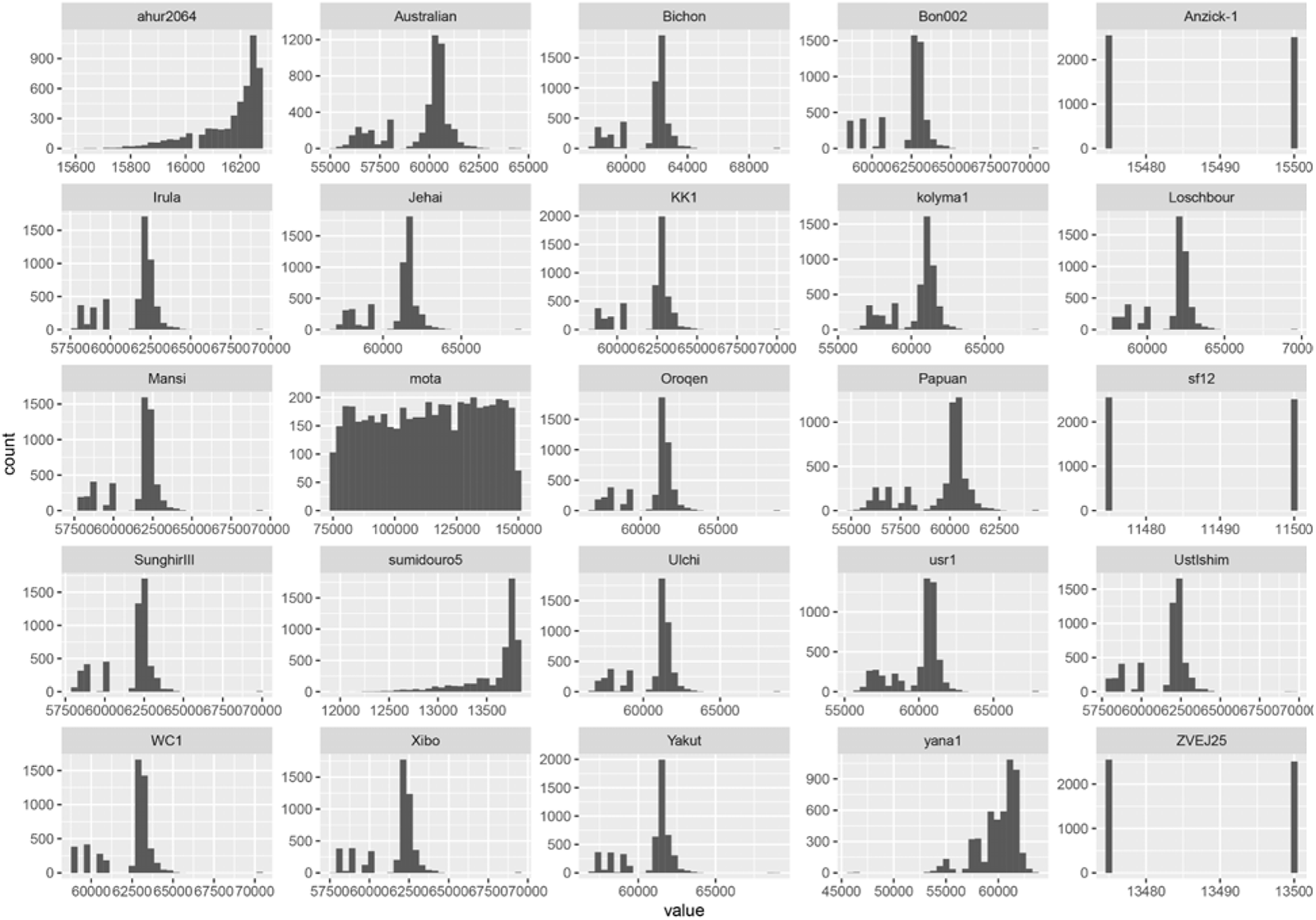
Histogram of earliest arrival time per population for the simulations retained during the parameter estimation (5058).

**Extended Data Fig. 6.**
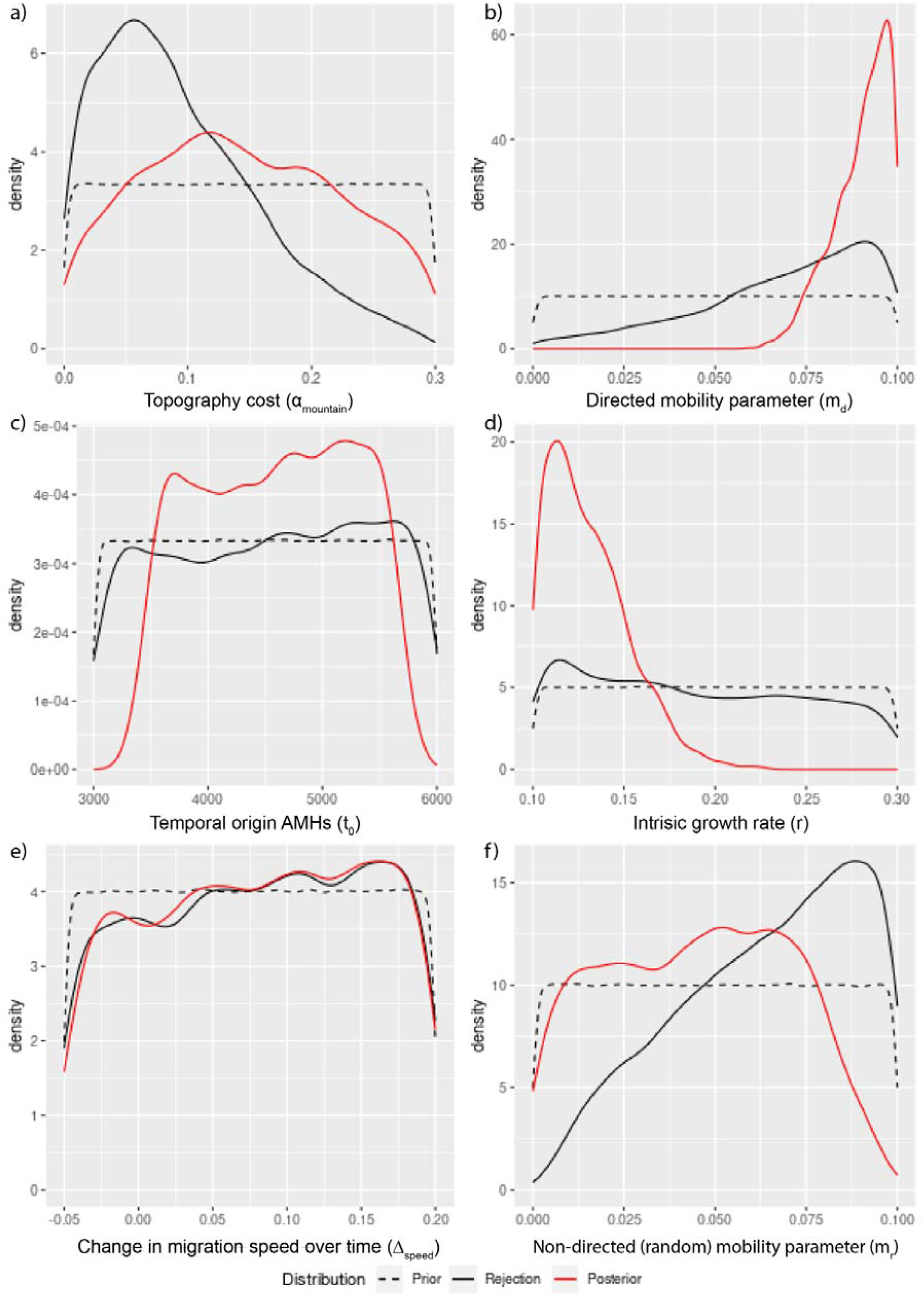
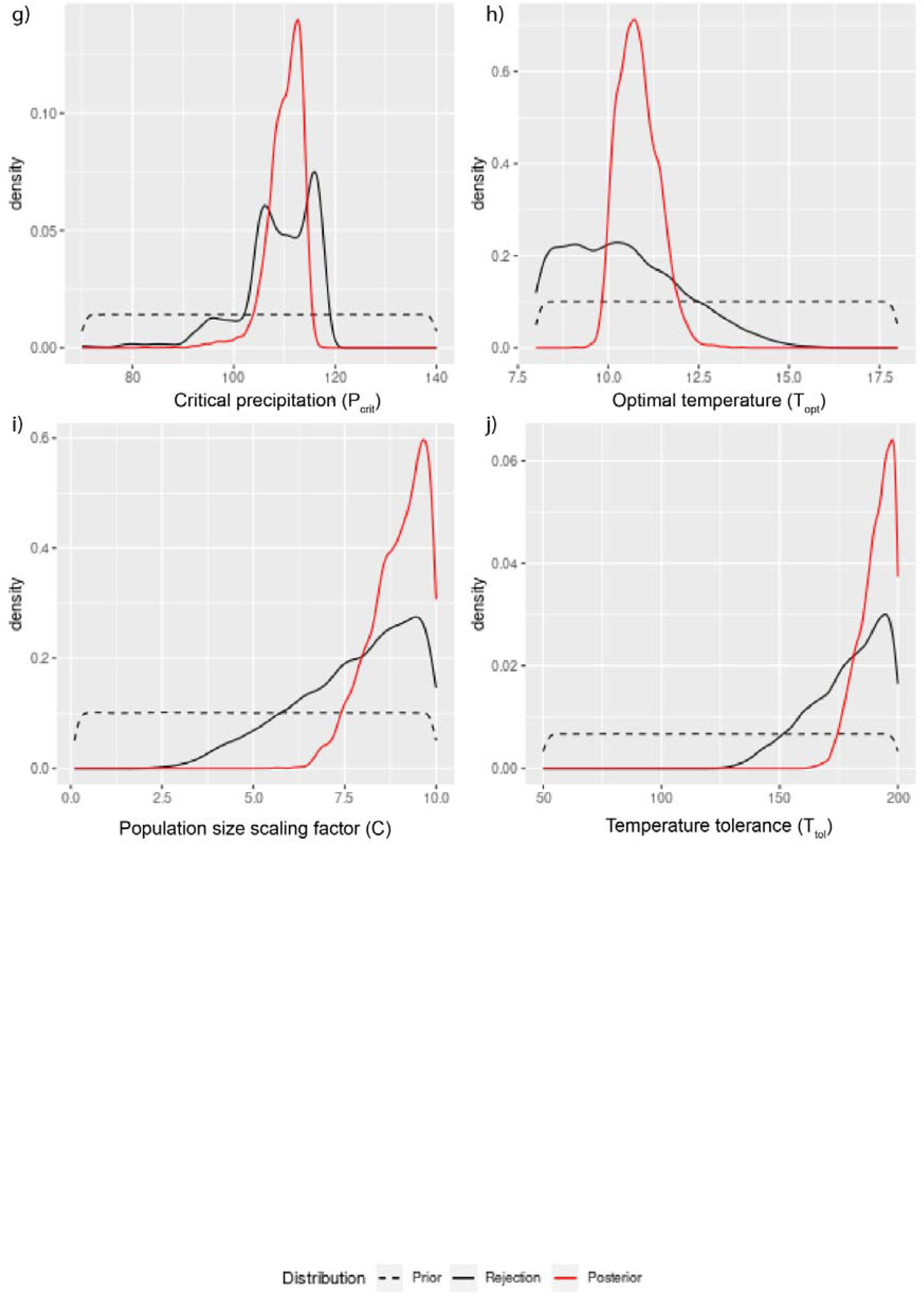
Posterior distributions for the parameter estimation in an ABC framework. Local linear regression, rejection approach and prior distributions are represented by the red, black and dashed black line respectively.

**Extended Data Fig. 7.**
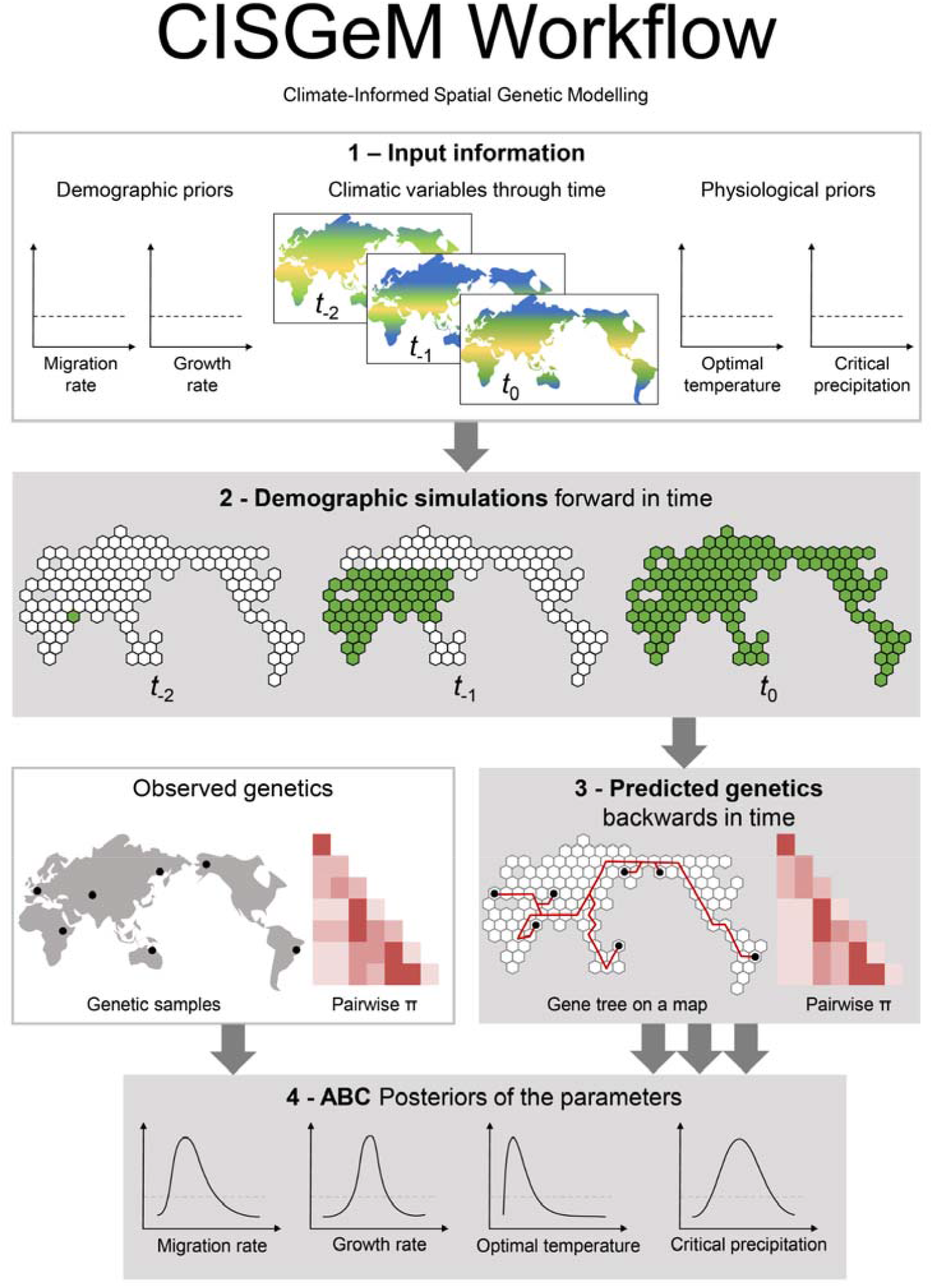
CISGeM workflow. CISGeM relies on **1)** a set of input information which includes prior distributions for both demographic and physiological parameters, and paleoclimate reconstructions for the period of interest. For every simulation, a value is picked at random from each prior distribution and **2)** the demography is generated using forward simulations. The availability of a cell (and therefore the resources to live on) is determined by the climatic variables through time combined with the appropriate demographic parameters. After the demography has been run from past to present, the genetics is traced backwards in time using a Discrete Time Wright Fisher model, and **3)** genetic quantities are predicted for the samples of interest. A large number of combinations of parameters are tested to cover exhaustively the prior distributions. Finally, **4)** the predicted genetics is compared against the observed genetics in an Approximate Bayesian Computation framework to produce posterior distributions for each parameter.

**Extended Data Fig. 8:**
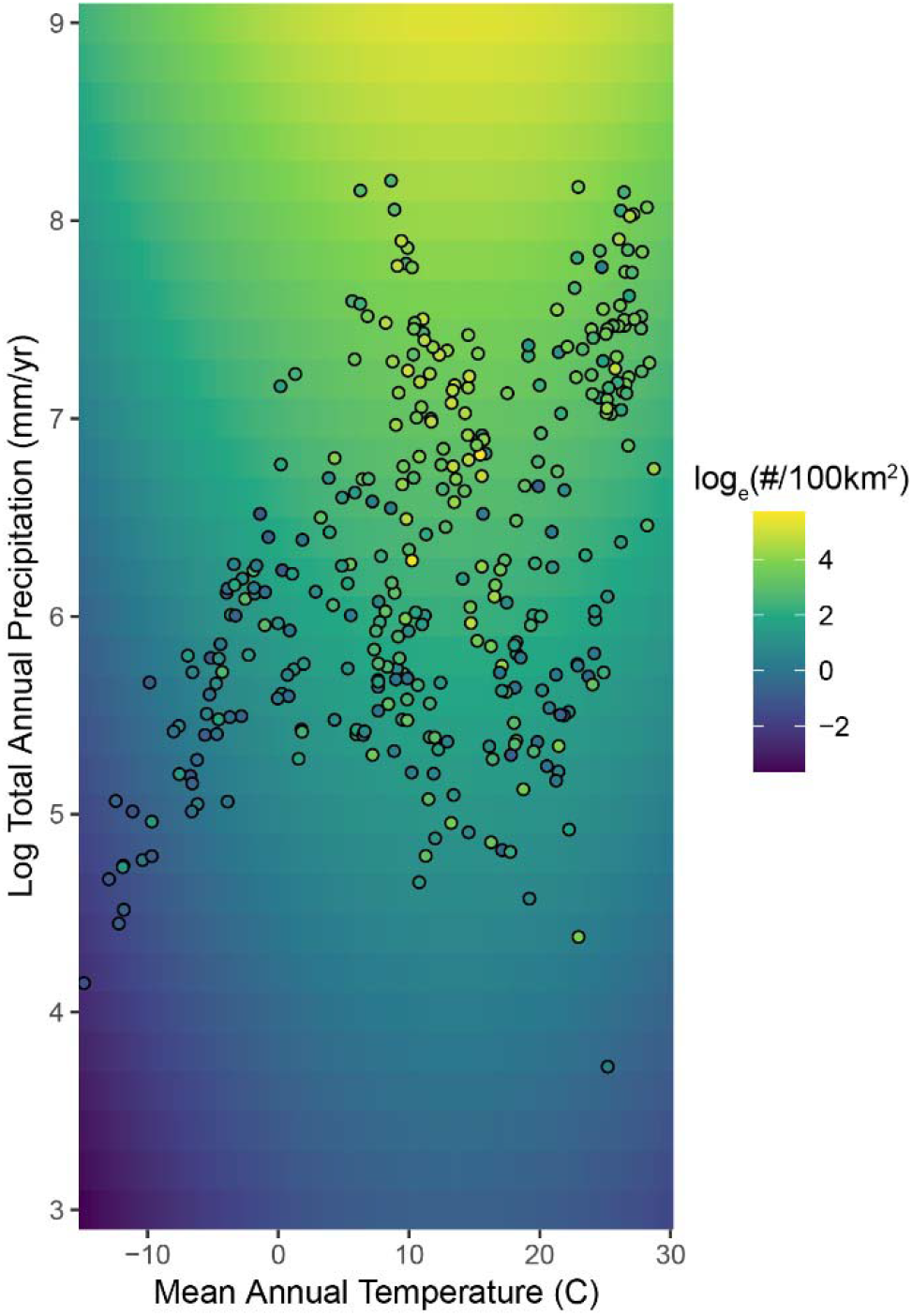
Effect of mean annual temperature and annual precipitation to census population sizes of modern hunter-gatherer groups. The relationship between mean annual temperature and annual precipitation (Pearson’s r: 0.47; p<0.001) provides a good fit to the census population sizes of modern hunter-gatherer populations.

**Extended Data Table 1** | metadata for ancient samples.

**Extended Data Table 2** | metadata for modern samples

**Extended Data Table 3** | List of parameters included in the demographic model.

**Extended Data Table 4** | Parameter estimation performed in an Approximate Bayesian Computation framework with local linear regression.

⍰ Upper and lower limits of the 95% credible interval.

⍰ Uniform probability, in the range of the two values.

⍰ Generation time 25 year.

