## Supplementary Information for "Climate and mountains shaped human ancestral genetic lineages"

### 1. Genetic summary statistics

#### Neutral loci

We selected 9178 loci which were 10kbp in length along the genome to represent neutral genetic variation. Filters excluding problematic regions or regions that may be under selection were applied to the human genome. Excluded regions include coding regions, conserved elements, recombination hotspots (regions with recombination rates >10cM/Mb), repetitive regions, and regions with poor mapping or sequencing quality. Filters used are defined in^1^ and kindly sent by Ilan Gronau. The total number of retained sites after this filtering has a maximum of 566,640,909 bp.

Contiguous intervals of 10kbp were chosen on these sites using a sliding window approach. Windows were retained if 7,500 sites or more were present, and a subset of these were selected with a minimum inter-locus distance of 50kbp. This minimum 50kbp distance between loci was chosen so that the chance of recombination was sufficiently high that loci could be treated as unlinked. This distance was determined by an approximate calculation similar to^2^. Mean recombination rate is assumed to be 10-8 per bp per generation, an average generation time of 25 years and a minimum average genomic divergence time of ~200,000 years. Following the logic in^2^, the expected number of recombinations in a 50kbp interval is at least 2 x 200,000 x 10-8/25 x 50000 = 8.

These loci were identified using 7 modern samples distributed worldwide: French (HGDP00526), Han (HGDP00783^3^), Yoruba (HGDP00928)^4^, Korean (KPGP-00288, KPGP-00347)^3^ and Singaporean Malay (SS6002734 and SS6002753)^5^. 109 of these loci were removed after filtering for coverage and CpG sites. In total, 9069 windows remained after filtering and they have been used in subsequent analyses.

#### Genetic diversity

The number of transitions (ts) and transversions (tv) was calculated for each sample within the HG_EXT dataset. The non UDG-treated samples (Bichon, WC1, Yana1, Kolyma1, Sumidouro5, Anzick-1 and Mota) show an excess of transitions because of DNA damage and degradation (see Supplementary Figure 1, panel a and c). All modern and ancient UDG-treated samples have similar ts/tv ratio except Loschbour which shows an unusually low ts/tv ratio. The mean ts/tv ratio (1.7245, red line in Supplementary Figure 1, panel b) has been calculated as average across all samples but the ancient non UDG-treated individuals and Loschbour.

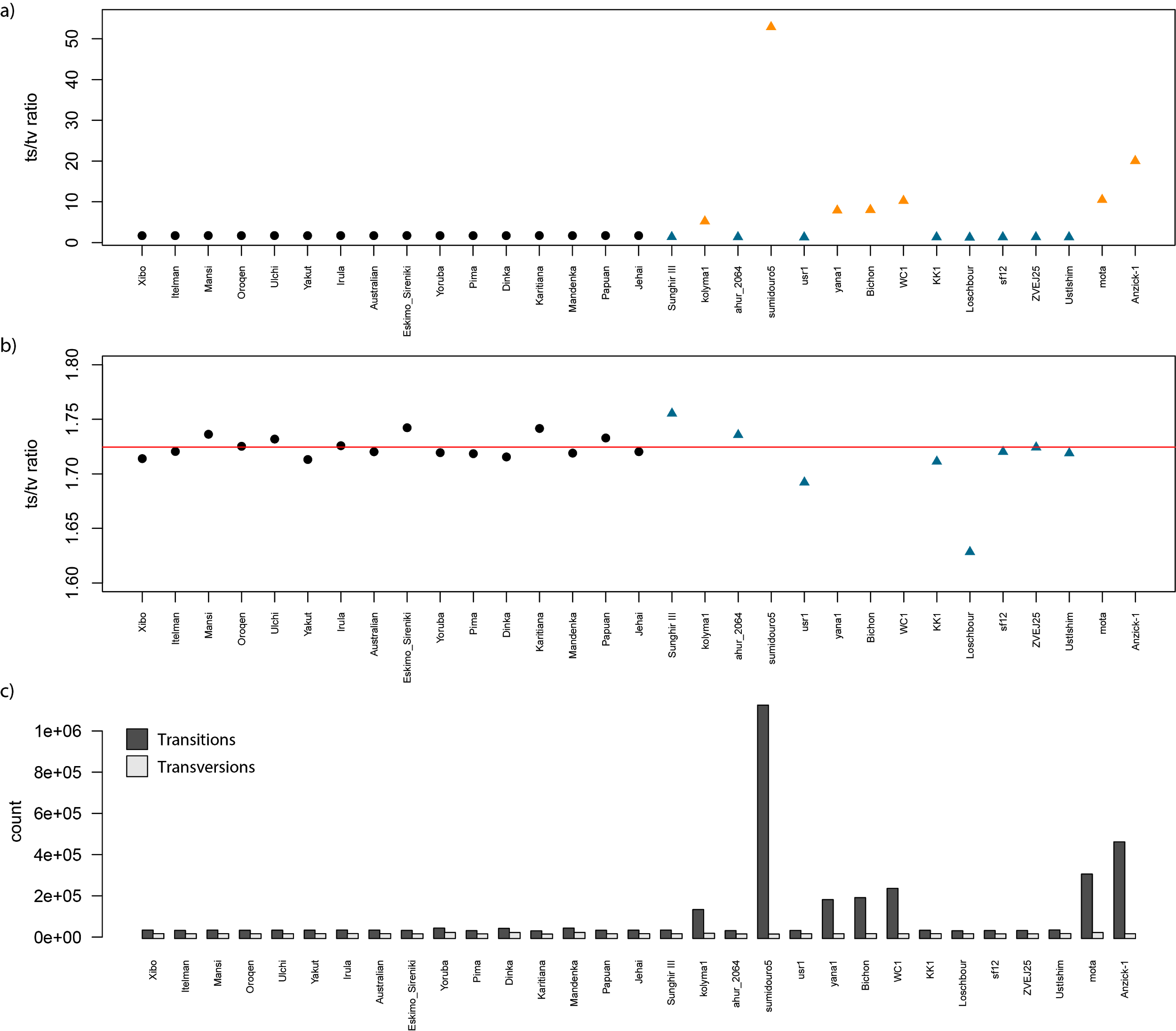

**Supplementary Figure 1**: a) Whole-genome transitions/transversions ratio (ts/tv) per sample based on 84,782,047 sites. Ancient and modern samples are represented by triangles and circles respectively. UDG and non-UDG treated samples are in blue and orange respectively. b) same as in a) but with a different y axis to focus on the ts/tv ratio among modern and UDG-treated ancient samples. Average ts/tv across all modern and ancient UDG-treated samples (except Loschbour) is represented by the red line. c) Number of transitions (ts) and transversions (tv) per sample.

For all samples except ancient non UDG-treated samples and Loschbour, pairwise π was calculated directly from whole-genome data (π_wg_). However, the same approach could not be used for the samples discarded above. Therefore, we calculated the number of pairwise differences on transversions only (diff_tv) and we used the average ts/tv ratio previously calculated to rescale the estimates of pairwise π based on transversion only to whole genome (π_wg_from_tv_) with the following formula: π_wg_from_tv_= (((diff_tv * average_ts/tv)+diff_tv) / total_sites). We also calculated the correlation between π_wg_ and π_wg_from_tv_ for all modern and ancient UDG-treated samples (except Loschbour) to assess how well the estimates based on transversions only represent whole-genome values. The correlation value is 0.997 and estimates were also plotted in Supplementary Figure 2.

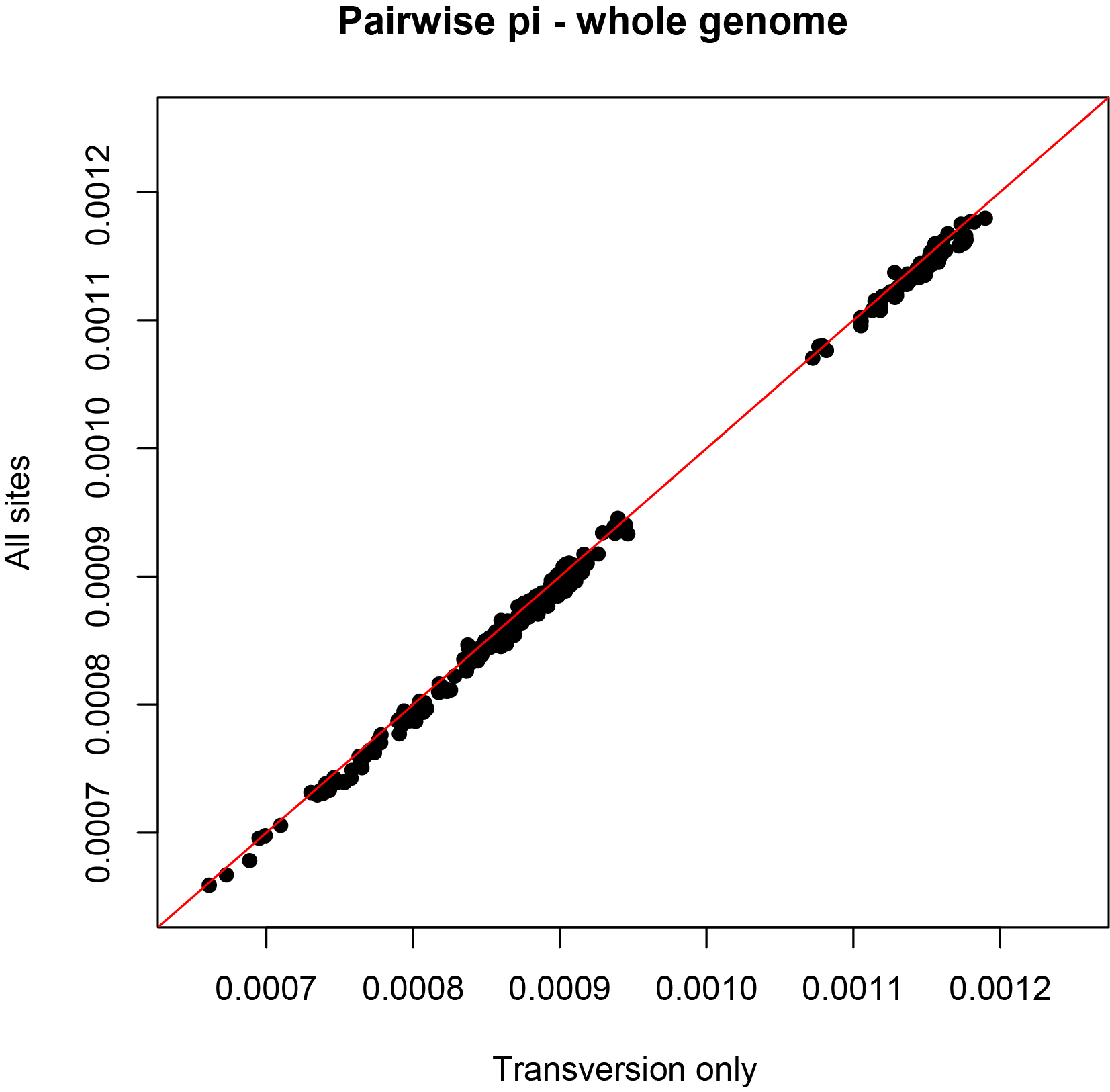

**Supplementary Figure 2**: Correlation between π_wg_ and π_wg_from_tv_.

The same approach was used for the HG_EXT dataset subset for the 9069 windows identified above (total sites retained 9,618,572). To assess how well the 9069 windows represent the genetic diversity at the whole genome level, a correlation coefficient was computed between π_wg_ and π_windows_. The correlation coefficient is 0.977 and pairwise π estimates are plotted in Supplementary Figure 3.

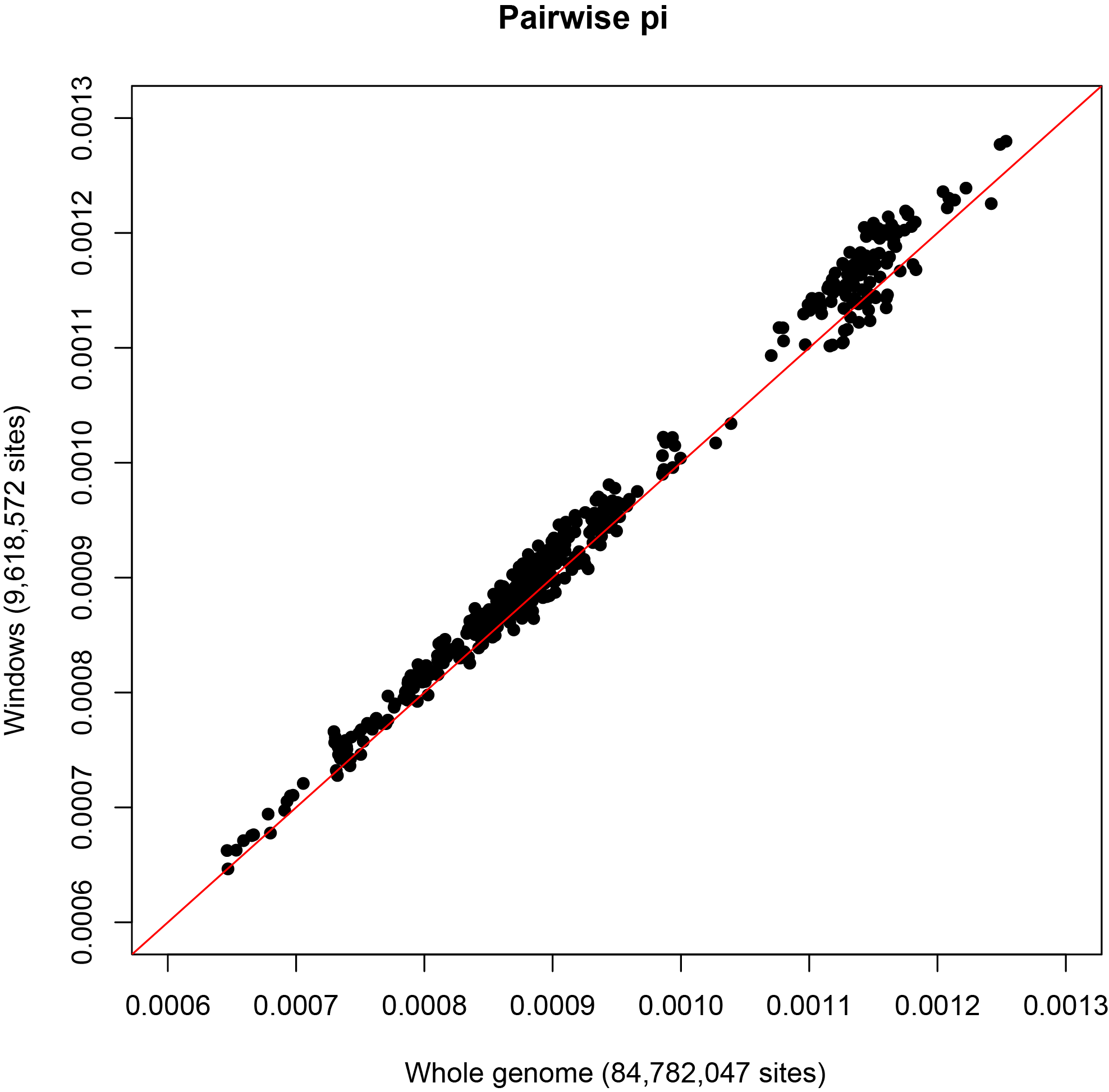

**Supplementary Figure 3**: Correlation between π_wg_ and π_windows_. All modern and ancient UDG-treated samples (except Loschbour) were included.

At this stage, the windows subset (9,618,572 sites) has been called as haploid in Bon002 with pileupCaller^6^ because the coverage did not allow calling diploid genotypes reliably. When merging Bon002, 494,784 sites were discarded because of missing data bringing the total number of sites to 9,123,788. We also checked for the presence of tri-allelic sites, but none were found. The merged VCF file includes 25 samples (16 ancient and 9 modern individuals) and represents the genetic dataset used for the spatial simulations. We then converted the diploid vcf into a haploid vcf to match the approach used in msprime^7,8^ where haploid chromosomes are simulated. The number of transitions and transversions has been recounted and the mean ts/tv ratio (1.7264, red line in Supplementary Figure 4, panel b) has been recalculated as average across all samples but the ancient non UDG-treated individuals and Loschbour. Estimates of pairwise π for non UDG-treated samples and Loschbour were calculated by rescaling the values based on transversions only as described above.

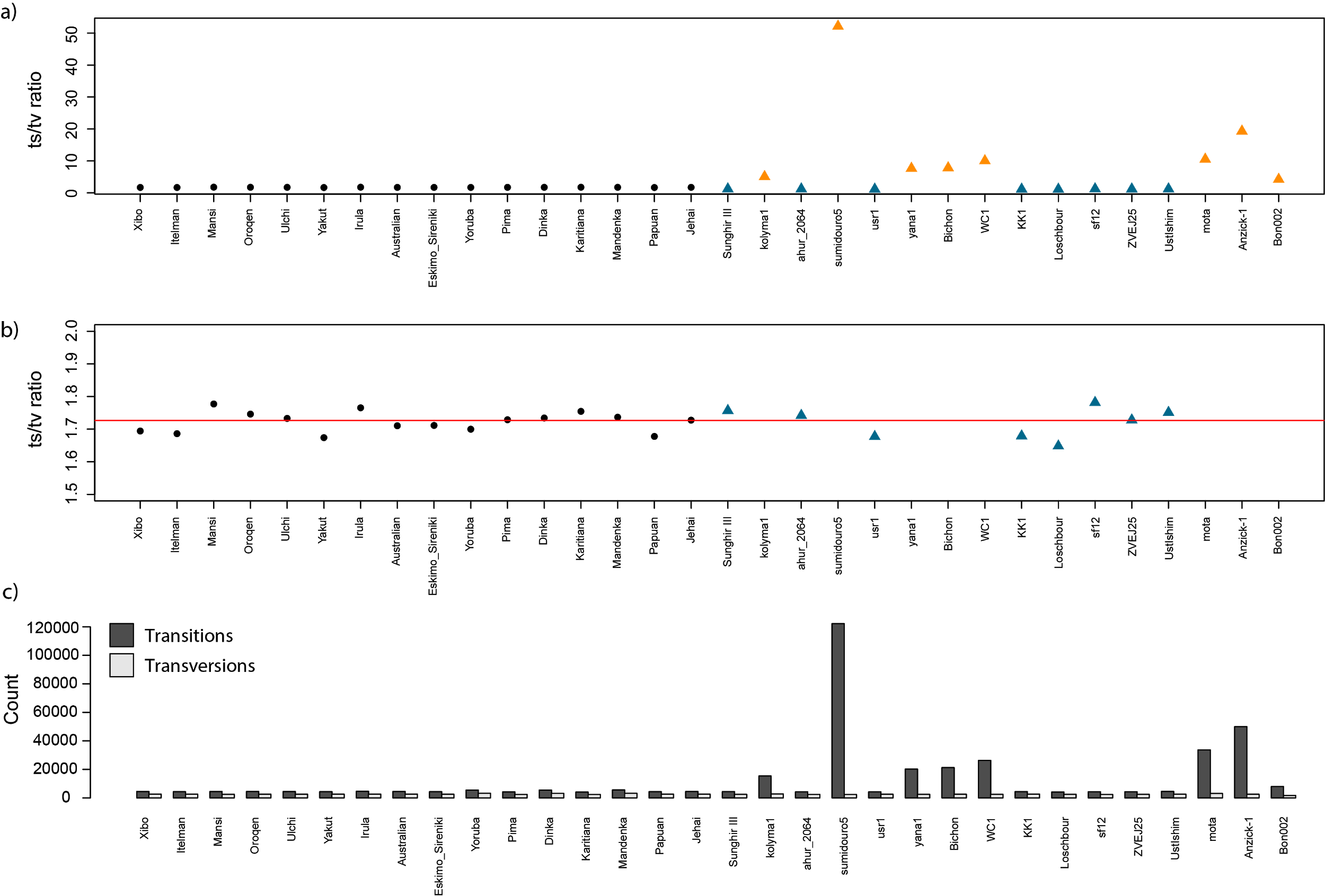

**Supplementary Figure 4**: a) Transitions/transversions ratio (ts/tv) per sample based on the windows subset after merging Bon002 (9,123,788 sites). Ancient and modern samples are represented by triangles and circles respectively. UDG and non-UDG treated samples are in blue and orange respectively. b) same as in a) but with a different y axis to focus on the ts/tv ratio among modern and UDG-treated ancient samples. Average ts/tv across all modern and ancient UDG-treated samples (except Loschbour) is represented by the red line. c) Number of transitions (ts) and transversions (tv) per sample.
